# Integrated Single-Cell Multiomic Profiling of Caudate Nucleus Suggests Key Mechanisms in Alcohol Use Disorder

**DOI:** 10.1101/2024.08.02.606355

**Authors:** Nick Green, Hongyu Gao, Xiaona Chu, Qiuyue Yuan, Patrick McGuire, Dongbing Lai, Guanglong Jiang, Xiaoling Xuei, Jill L. Reiter, Julia Stevens, Greg T. Sutherland, Alison M. Goate, Zhiping P. Pang, Paul A. Slesinger, Ronald P. Hart, Jay A. Tischfield, Arpana Agrawal, Yue Wang, Zhana Duren, Howard J. Edenberg, Yunlong Liu

**Affiliations:** Indiana University School of Medicine, Department of Medical and Molecular Genetics, Indianapolis, Indiana 46202, United States; Center for Computational Biology and Bioinformatics, Indiana University School of Medicine, Indianapolis, Indiana 46202, United States; Clemson University, Department of Genetics and Biochemistry, Greenwood, SC 29646, United States; New South Wales Brain Tissue Research Centre, Charles Perkins Centre and School of Medical Sciences, Faculty of Medicine and Health, The University of Sydney, Camperdown, NSW 2006, Australia; Nash Family Department of Neuroscience, Icahn School of Medicine at Mount Sinai, New York, New York 10029, United States; Department of Genetics & Genomic Sciences, Icahn School of Medicine at Mount Sinai, New York, New York 10029, United States; Human Genetics Institute, Rutgers University, Piscataway, New Jersey 08854, United States; Department of Neuroscience and Cell Biology and The Child Health Institute of New Jersey, Rutgers Robert Wood Johnson Medical School, New Brunswick, New Jersey 08901, United States; Department of Cell Biology & Neuroscience, Rutgers University, Piscataway, New Jersey 08854, United States; Department of Genetics, Rutgers University, Piscataway, New Jersey 08854, United States; Department of Psychiatry, Washington University School of Medicine in St. Louis, St. Louis, Missouri 64110, United States; Department of Biochemistry and Molecular Biology, Indiana University School of Medicine, Indianapolis, Indiana 46202, United States

## Abstract

Alcohol use disorder (AUD) induces complex transcriptional and regulatory changes across multiple brain regions including the caudate nucleus, which remains understudied. Using paired single-nucleus RNA-seq and ATAC-seq on caudate samples from 143 human postmortem brains, including 74 with AUD, we identified 17 distinct cell types. We found that a significant portion of the alcohol-induced changes in gene expression occurred through altered chromatin accessibility. Notably, we identified novel transcriptional and chromatin accessibility differences in medium spiny neurons, impacting pathways such as RNA metabolism and immune response. A small cluster of D1/D2 hybrid neurons showed distinct differences, suggesting a unique role in AUD. Microglia exhibited distinct activation states deviating from classical M1/M2 designations, and astrocytes entered a reactive state partially regulated by *JUND*, affecting glutamatergic synapse pathways. Oligodendrocyte dysregulation, driven in part by *OLIG2*, was linked to demyelination and increased TGF-β1 signaling from microglia and astrocytes. We also observed increased microglia-astrocyte communication via the IL-1β pathway. Leveraging our multiomic data, we performed cell type-specific expression quantitative trait loci analysis, integrating that with public genome-wide association studies to identify AUD risk genes such as *ADAL* and *PPP2R3C*, providing a direct link between genetic variants, chromatin accessibility, and gene expression in AUD. These findings not only provide new insights into the genetic and cellular mechanisms in the caudate related to AUD but also demonstrate the broader utility of large-scale multiomic studies in uncovering complex gene regulation across diverse cell types, which has implications beyond the substance use field.

## Introduction

Excessive alcohol use creates many serious physical, emotional, and social problems and is responsible for about 3 million deaths worldwide each year.^1^ Many deaths in the United States result from alcohol use disorder (AUD) (nccd.cdc.gov/DPH_ARDI). AUD is a serious and common psychiatric disorder that is characterized by excessive alcohol consumption and consequent psychological and interpersonal problems stemming from preoccupation with and a loss of control over drinking.^2^ The risk of developing alcohol use disorder (AUD) depends on both genetic and environmental factors. While recent large-scale genome-wide association studies (GWAS) have identified hundreds of variants associated with alcohol consumption^3,4^ and AUD^5-7^ it is not yet clear how these variants contribute to AUD.

Beyond these predisposing differences in the genome, AUD is likely associated with dynamic alterations in gene expression and chromatin conformation, plausibly in brain regions associated with onset and maintenance of motivated and rewarding behaviors, stress responsivity and cognitive control. Early studies using microarray analysis to study the effects of chronic ethanol consumption in rats found significant changes in expression in genes in several brain regions, including the nucleus accumbens,^8,9^ extended amygdala,^10,11^ and ventral tegmental area,^12^ and studies in human lymphoblastoid cell lines and postmortem tissue identified expression changes associated with alcohol dependence.^13,14^ Subsequent bulk RNA-sequencing (RNA-seq) studies have uncovered differentially expressed genes in several brain regions, including the rat hippocampus, prefrontal cortex^15^, raphe nuclei,^16^ and periaqueductal gray^17^, in human lymphoblastoid^18^ and neuroblastoma^19^ cell lines, and in human postmortem tissue from the hippocampus,^20^ prefrontal cortex,^21,22^ and striatum.^23^

However, the individual brain regions contain a diversity of cell types that may bear unique transcriptional signatures that cannot be detected in bulk RNA sequencing data even with computational deconvolution techniques. Single-cell/single-nucleus RNA sequencing has enabled measurement of the distribution and characterization of different cell types in a tissue sample and of gene expression in each of these individual cells. An early single-nucleus RNA sequencing (snRNA-seq) study examined gene expression in nuclei from the prefrontal cortex of individuals with and without AUD.^24^ Despite a small sample size (7 individuals), they identified seven major cortical cell types and found differences in expression associated with AUD within six cell types, particularly in astrocytes, oligodendrocytes and microglia, some in neuroinflammation-related genes.

Changes in gene expression may have manifold etiologies. One likely epigenetic precursor is the accessibility of chromatin, which exerts a *cis*-regulatory effect on gene expression. For example, a recent study using an assay for transposase-accessible chromatin with sequencing (ATAC-seq^25^) found differences in chromatin accessibility associated with chronic and acute alcohol exposure in the rat amygdala.^26^ However, AUD-associated linkages between open chromatin regions and gene expression are likely to be regionally and cell-type specific. The advent of single nucleus multiome experiments (sn-multiome), assaying both chromatin accessibility and gene expression within the same cell, provides remarkable opportunities to draw causal inferences regarding mechanisms underlying AUD-associated gene expression.

The caudate nucleus forms part of the dorsal striatum (and more broadly, the basal ganglia), a key component of the executive control loop that is recruited in the onset and maintenance of AUD.^27^ The caudate has been implicated in cue-elicited activation, dopamine increase, and in subjective reports of craving.^28,29^ In animal models, chronic ethanol exposure alters neural circuits in the basal ganglia^30,31^ with a recent study reporting differences in gene expression in the dorsal striatum of alcohol-preferring rats^32^. A transcription-wide association study found that among 13 human brain tissues, the caudate was the region with the most genes whose predicted expression was associated with problematic alcohol use (PAU), a trait that combines AUD with problematic alcohol drinking.^5^ However, the caudate harbors multiple cell-types^33^ and cell-type-specific characterization of the AUD-associated transcriptome in the human caudate is lacking.

To meet this gap in knowledge, we sought to provide a comprehensive view of AUD-related differences in gene expression and chromatin accessibility in specific cell types within the human caudate nucleus and infer mechanisms underlying these changes. We performed a high-throughput snRNA-seq experiment on human postmortem samples from the caudate nucleus of 143 donors, 74 with and 69 without AUD, obtaining transcriptomic data from over 1.1 million cells. To compare the transcriptome with the open chromatin status of the same cells, we also performed an sn-multiome analysis from these same caudate samples. Sn-multiome experiments have been used in recent years to study human brain development^34,35^ and neuropsychiatric diseases such as Alzheimer’s disease,^36,37^ but these studies have been limited by small sample sizes or shallow sequencing approaches, or studies in which sn-multiome data were supplemented with single-cell data generated in separate datasets.^38^ Here, we performed both a sn-multiome and a deep high-throughput snRNA-seq experiment in the same large sample cohort, allowing us to both robustly identify rare cell types and measure small differences in both gene expression and chromatin accessibility in the same nuclei.

We profiled AUD-associated differences in gene expression and chromatin accessibility in different cell types, the biological pathways underlying these differences, and AUD-associated differences in transcription factor activity and cell-cell communication in major glial cell types (Fig. 1). This study provides a comprehensive profile of AUD-related differences in the caudate nucleus and identifies potential mechanisms of AUD and novel directions for further exploration. Our novel experimental approach highlights the broader utility of large-scale multiomic studies for identifying regulatory mechanisms, which can be applied to other neurological and psychiatric conditions.

**Figure 1:**
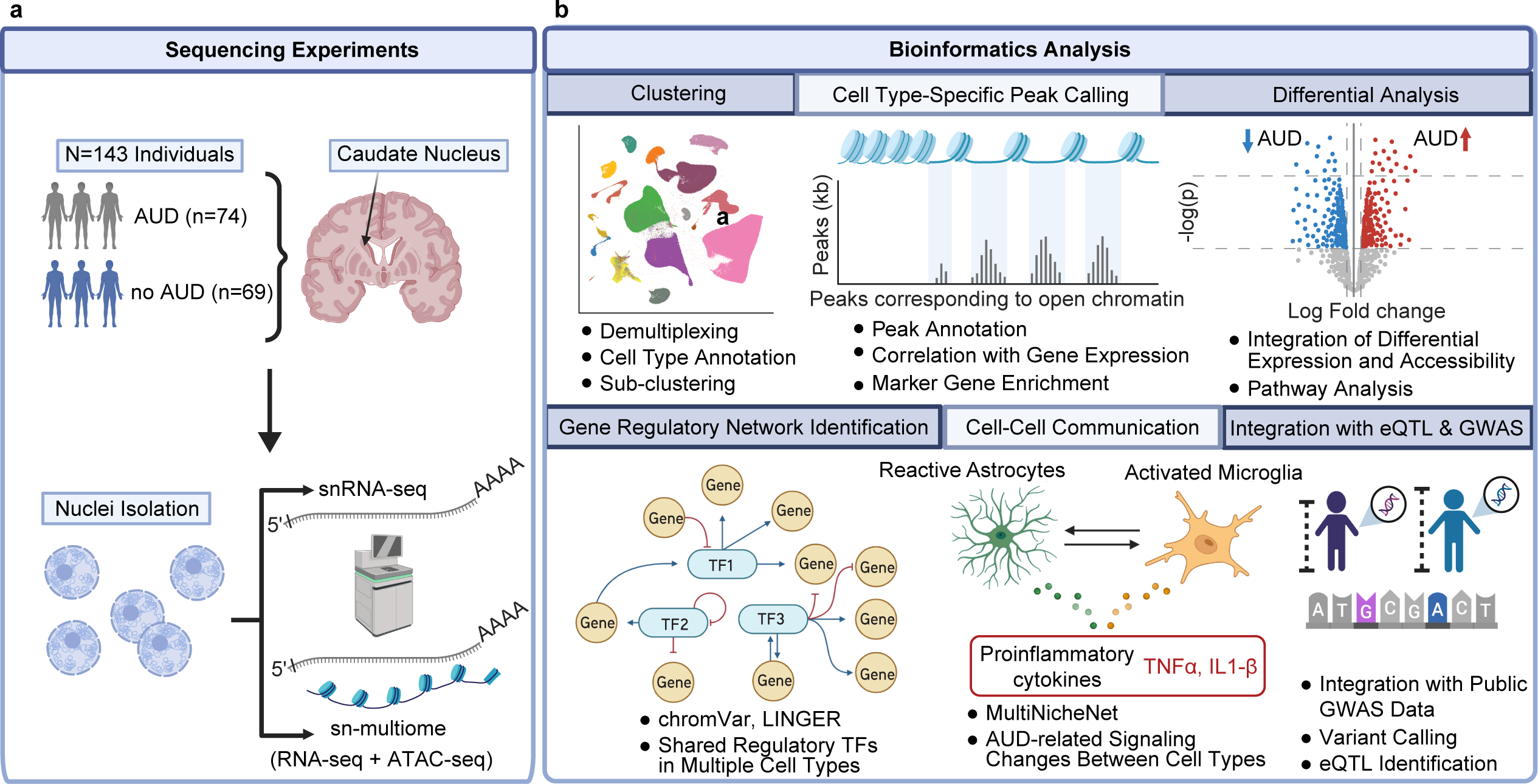
Experimental and bioinformatics workflow. **a,** Sequencing experiments included single-nucleus RNA-seq (snRNA-seq) and single-nucleus multiome (sn-multiome) of 163 postmortem brain samples from individuals with and without alcohol use disorder (AUD). **b,** Bioinformatics approaches and tools used for analysis. See Online Methods for details.

## Results

### Characterization of 17 Major Cell Types in the Caudate Nucleus

Samples from the caudate nucleus of post-mortem brains from the New South Wales Brain Tissue Resource Centre at the University of Sydney were sequenced in the sn-multiome assay, in which transcription levels and chromatin accessibility were measured in the same nuclei; most were also sequenced using the 10X HT snRNA-seq assay. After demultiplexing and data processing, samples with < 200 cell barcodes were removed, leaving 163 samples – 82 with and 81 without AUD; 128 male and 34 female. Low quality nuclei were filtered out from further analyses based on number of genes, number of molecules, and percentage of mitochondrial DNA (see ‘Initial Quality Control’ in Online Methods), leaving gene expression levels for 1,307,323 nuclei and chromatin accessibility (ATAC-seq) for 267,100 of these nuclei (Demographics are in Supplementary Tables 1-2, and detailed experimental procedures are in Online Methods). Graph-based clustering of the snRNA-seq data of the 163 samples identified 17 distinct cell clusters (Fig. 2A, B). Three subtypes of medium spiny neurons (MSNs, the GABAergic projection neurons of the striatum) were identified: D1-type MSNs, D2-type MSNs, and a third subtype marked by both *DRD1* and *DRD2* expression (D1/D2 neurons). Four small populations of GABAergic interneurons were identified, including parvalbumin-expressing fast-spiking (FS), neuropeptide Y/somatostatin/nitric oxide synthase-expressing low threshold-spiking (LTS), calretinin-expressing (CR), and cholecystokinin-expressing (CCK). A small cluster of cholinergic neurons was also identified (Ach). In addition to neurons, several glial cell populations were observed, including oligodendrocytes (the most prevalent cell type; 28.2% of the nuclei), oligodendrocyte progenitor cells (OPCs), astrocytes, ependymal cells, and microglia. Other cell types included non-microglial macrophages, endothelial cells, and vascular smooth muscle cells.

**Figure 2:**
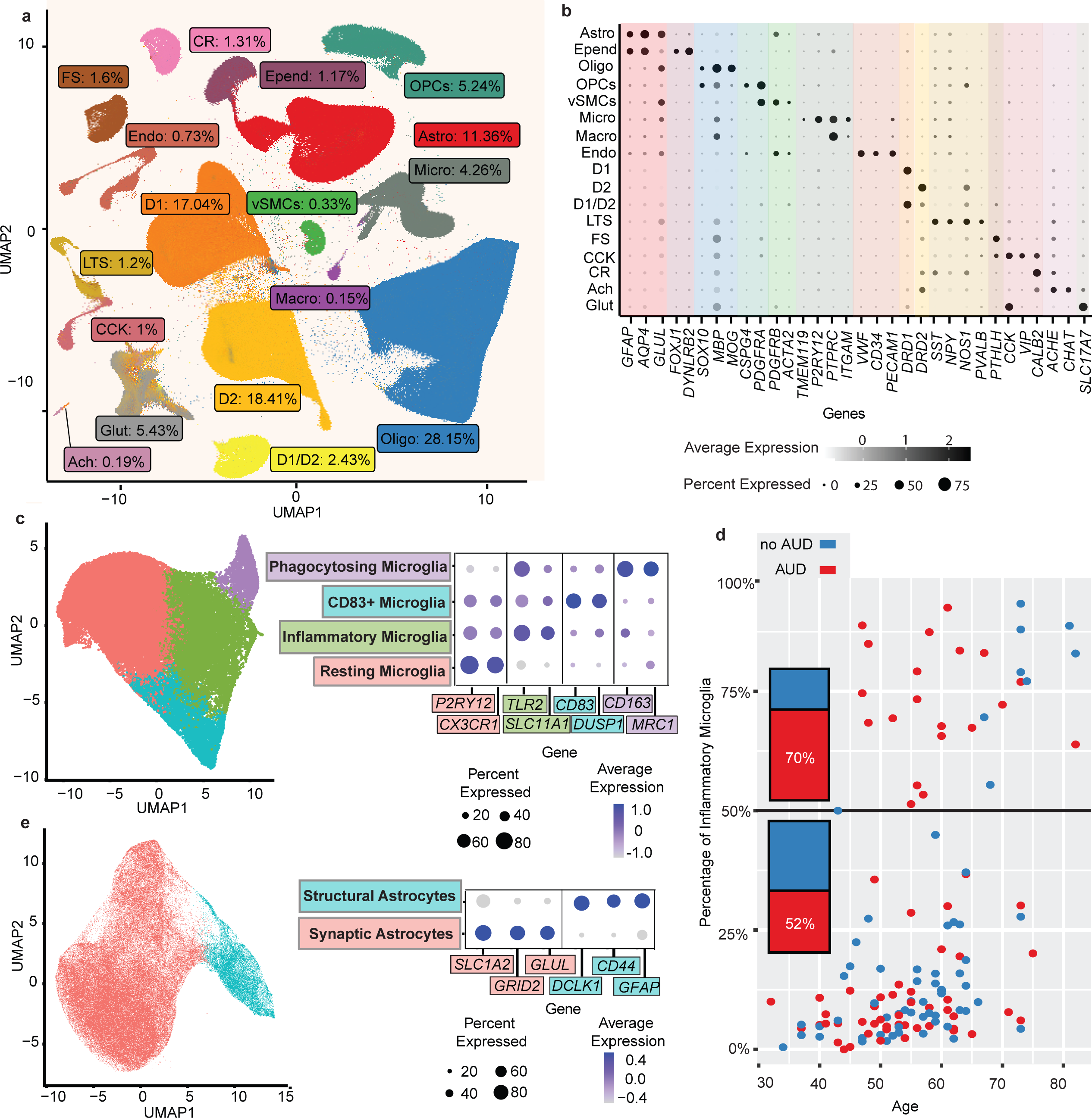
Cell type landscape of the caudate nucleus in alcohol use disorder. **a,** UMAP plot of the 1,307,323 nuclei profiled in the snRNA-seq and sn-multiome assays; visualization shown is based on the snRNA-seq profile. Nuclei are labeled by cell type and cell type proportion among all snRNA-seq cells. Cell types: cholinergic neurons (Ach), astrocytes (Astro), cholecystokinin-expressing interneurons (CCK), calretinin-expressing interneurons (CR), D1-type medium spiny neurons (D1), D2-type medium spiny neuron (D2), medium spiny neurons expressing both D1 and D2 receptors (D1/D2), endothelial cells (Endo), ependymal cells (Epend), fast-spiking interneurons (FS), glutamatergic neurons (Glut), low-threshold spiking interneurons (LTS), non-microglial macrophages (Macro), microglia (Micro), oligodendrocytes (Oligo), oligodendrocyte progenitor cells (OPCs), and vascular smooth muscle cells (vSMCs). **b,** Normalized expression in each cell type of the marker genes used to identify cell types. Dot size corresponds to the percentage of cells expressing the gene; dot intensity indicates average gene expression level. **c,** Left, UMAP of microglial cells, colored by subcluster. Right, dotpot of expression and prevalence of representative marker genes for each microglial subcluster. **d,** Scatter plot showing the proportion of inflammatory microglia in each sample and the subject’s age (red, AUD; blue, control). Bars on the left quantify the ratio of individuals with AUD to without AUD among those with >= 50% (above) or < 50% (below the solid black line) of microglia in the inflammatory state. **e,** Left, UMAP of astrocyte cells, colored by subcluster. Right, Dotpot of expression and prevalence of representative marker genes for each subcluster.

Unexpectedly, glutamatergic neurons were found, although the caudate is known not to contain cell bodies of excitatory neurons. Their presence could reflect inadvertent inclusion of another brain region at the time of dissection; therefore, twenty samples that contained >10% of glutamatergic neurons were removed from all subsequent analyses (Supplementary Tables 1, 2), leaving 143 samples (74 with AUD, 69 without; 115 male, 28 female) with gene expression data for 1,121,762 nuclei, 250,537 of which also had ATAC data. There was no significant difference in relative abundance of cell types between samples from individuals with and without AUD (Extended Data Figure 1).

Graph-based subclustering was performed for several cell types separately, using data from the 143 samples. There were four subclusters of microglia, which roughly correspond to different states of microglial activation (Fig. 2C, Extended Data Fig. 2, Supplementary Tables 3-4). Subcluster 1 (“Resting Microglia”) uniquely expressed genes specific to quiescent microglia, such as *P2RY12* and *CX3CR1,* and was enriched for pathways relating to microglia migration. Subclusters 2 and 3 were both enriched for immune response-related genes, with subcluster 2 (“Inflammatory Microglia”) highly expressing genes involved in inflammation, such as *TLR2*. Subcluster 3 (“CD83+ Microglia”) was enriched for genes governing microglia activation, such as *CD83.*^39^ Subcluster 4 (“Phagocytosing Microglia”) was marked by high expression of genes involved in endocytosis and phagocytosis. There was a significant increase in the mean proportion of “Inflammatory Microglia” (subcluster 2) in individuals with AUD: 31%, as opposed to 23% in those without AUD (adjusted *p* value (*padj*) = 0.027). Also, 70% of individuals that had at least half of their microglia cells in the inflammatory state (subcluster 2) had AUD, while only 52% of individuals who had below half of their microglia in the inflammatory state had AUD (Fig. 2D).

A large astrocyte subcluster (“Synaptic Astrocytes”) was marked by higher expression of excitatory amino acid transporters 1 and 2 (glutamate transporters), glutamate receptor 2 (an AMPA receptor subunit), and glutamate synthase, suggesting that these astrocytes may play a role in maintaining glutamatergic synapses (Fig. 2E, Supplementary Tables 5-6). The other subcluster (“Structural Astrocytes”) was marked by high expression of cytoskeleton-related protein-coding genes *GFAP* and *DCLK1*, extracellular matrix protein tenascin C, and *CD44*, coding for a protein involved in cell adhesion and migration, and thus may be involved in structural support or tissue repair.

Finally, there were two subclusters of both D1 and D2-type neurons, representing matrix and striosome compartments,^40^ based on expression of genes specific to either the matrix or striosome regions of the striatum^41^ (Extended Data Fig. 2-3). 80% of D1 neurons and 83% of D2 neurons were within the matrix compartment, which makes up approximately 85% of the striatum.^40^ There was no significant difference in subcluster proportion by AUD status in either astrocytes or D1 and D2-type neurons.

### AUD-Associated Differences in Gene Expression

We performed differential gene expression analyses in thirteen major cell types in which there were greater than 50 cells of that type in more than 10 individuals with and 10 without AUD. We utilized a pseudobulk approach, summing counts across cells within each sample for each cell type, with sex, age, and ethnic origin as covariates. Samples were removed on a cell type-specific basis if the sample contained less than 50 cells of that cell type (See Supplementary Table 7 for a summary of the number of pseudobulk samples created for each cell type).

Eight cell types each contained over 700 differentially expressed genes (DEGs) (*padj* < 0.2) (Fig. 3A, Supplementary Table 7). We identified many changes in astrocytes, microglia, and in neurons, especially D1 and D2 medium spiny neurons. The magnitude of gene expression changes in neurons was generally smaller than in the astrocytes and microglia. In each of these cell types, more genes had higher expression in individuals with AUD than had lower. Many of the DEGs were differentially expressed in multiple cell types. Notably, astrocytes and oligodendrocytes had 833 DEGs in common, and D1 and D2 neurons had 538 DEGs in common (Extended Data Fig. 4). The differences in gene expression within D1 neurons and D2 neurons were highly correlated with each other (Pearson correlation = 0.83), while differences within other neuronal and non-neuronal cell types were more weakly correlated (Pearson correlation ranges = 0.10-0.46 for neuronal cell types and 0.05-0.27 for non-neuronal cell types; Fig. 3B, Supplementary Table 8). Interestingly, expression differences within D1/D2 neurons were much less correlated with either D1 or D2 neurons (*r =* 0.41 and 0.39 respectively).

**Figure 3:**
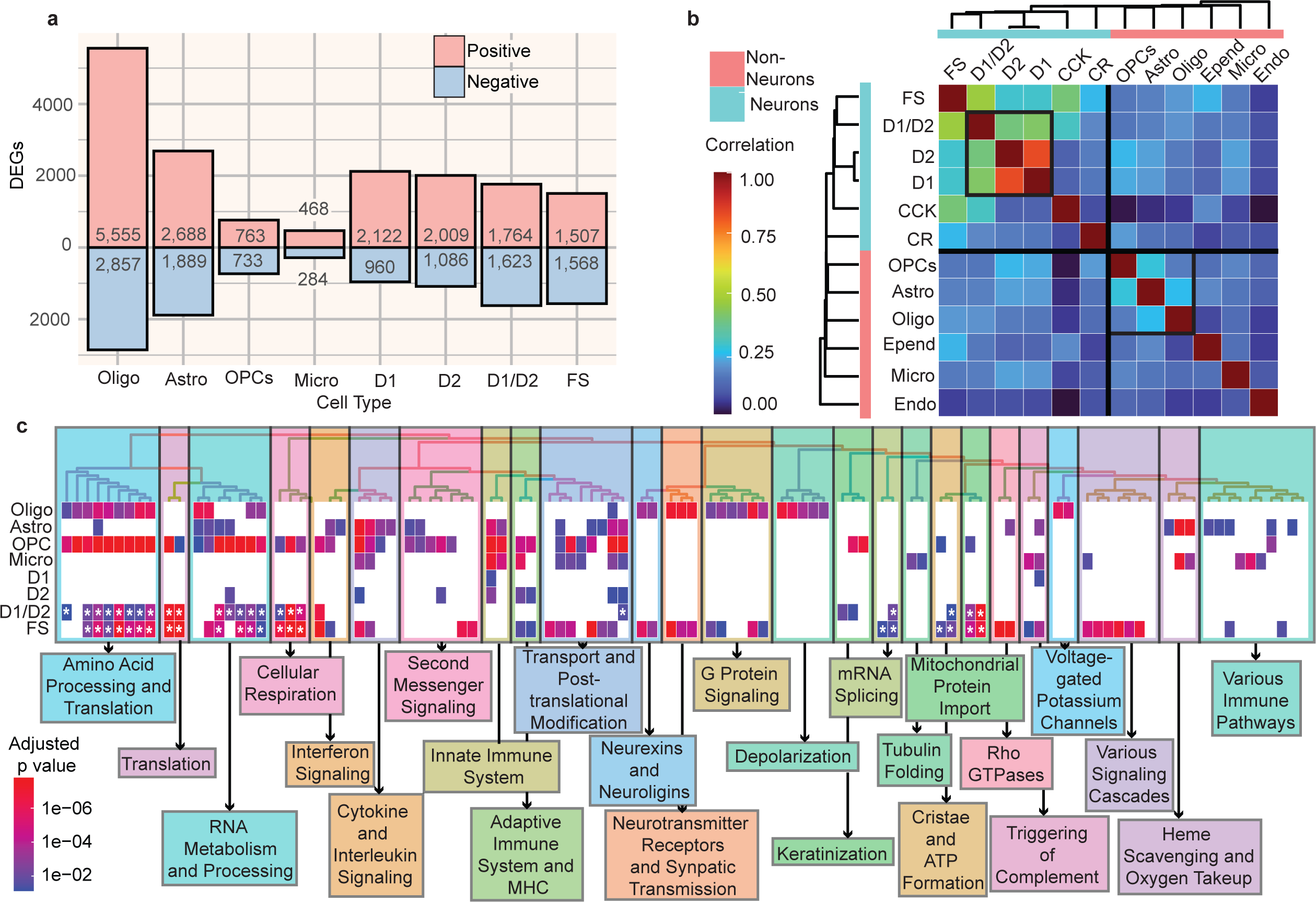
Characterization of AUD-associated changes in gene expression in the caudate nucleus. **a,** Barplot showing number of genes differentially expressed in individuals with AUD for the eight cell types which have over 100 differentially expressed genes. Red and blue indicate positively and negatively differentially expressed genes, respectively. See Fig. 2A caption for cell type abbreviations. **b,** Heatmap of Pearson correlation of gene expression changes (log2 fold changes) between cell types, hierarchically clustered by Pearson correlation. Black-outlined squares indicate groups of cell types with moderate correlation, namely, D1, D2, and D1/D2 neurons, and OPCs, astrocytes, and oligodendrocytes. **c,** Heatmap of biological pathways from the Reactome database enriched in brain samples from individuals with AUD in each cell type. The top 100 enriched pathways (based on smallest Benjamini-Hochberg adjusted p values) across all cell types are shown and were hierarchically clustered based on the number of genes shared between the pathways. Heatmap cell color indicates adjusted p value. Asterisk indicates negative enrichment score; all other pathways have positive enrichment scores. The pathways are divided into 25 clusters, which are manually labeled with a summary of pathways making up that cluster.

Gene set enrichment analysis using pathways from the Reactome database^42^ showed that genes that differ in expression with respect to AUD were enriched in hundreds of pathways in many cell types (Fig. 3C, Supplementary Table 9). Successive hierarchical and manual grouping of pathways revealed that many immune response pathways – such as the adaptive immune system, innate immune system, and cytokine signaling in immune system – were enriched in multiple cell types from individuals with AUD. In individuals with AUD, DEGs in oligodendrocytes were enriched for several pathways associated with synaptic regulation and depolarization, such as “Neurotransmitter Receptors and Postsynaptic Signal Transmission” and “Voltage Gated Potassium Channels”. D1/D2 and FS neurons had decreases in gene expression within pathways related to translation and metabolism.

### AUD-Associated Differences in Chromatin Accessibility

For the 267,100 cells in which both gene expression and chromatin accessibility data were available (see Extended Data Fig. 5 for cell type distribution for snATAC-seq cells), the chromatin accessibility (combined reads for each cell type) showed a strong signal near the transcription start sites (TSS) of its respective marker genes of several cell type-specific genes (Extended Data Fig. 6). Thirty-six percent of all open chromatin regions were shared among neuronal and non-neuronal cell types, while 34% of the peaks were unique to neurons and 30% unique to non-neurons (Fig. 4A). D1 neurons and D2 neurons had very similar open chromatin regions (Jaccard index = 0.8) and were less similar to D1/D2 neurons (Jaccard index = 0.47). Astrocytes, oligodendrocytes, and OPCs had moderately similar open chromatin regions, with Jaccard indices of approximately 0.4 between these cell types (Fig. 4B, Supplementary Table 10).

**Figure 4:**
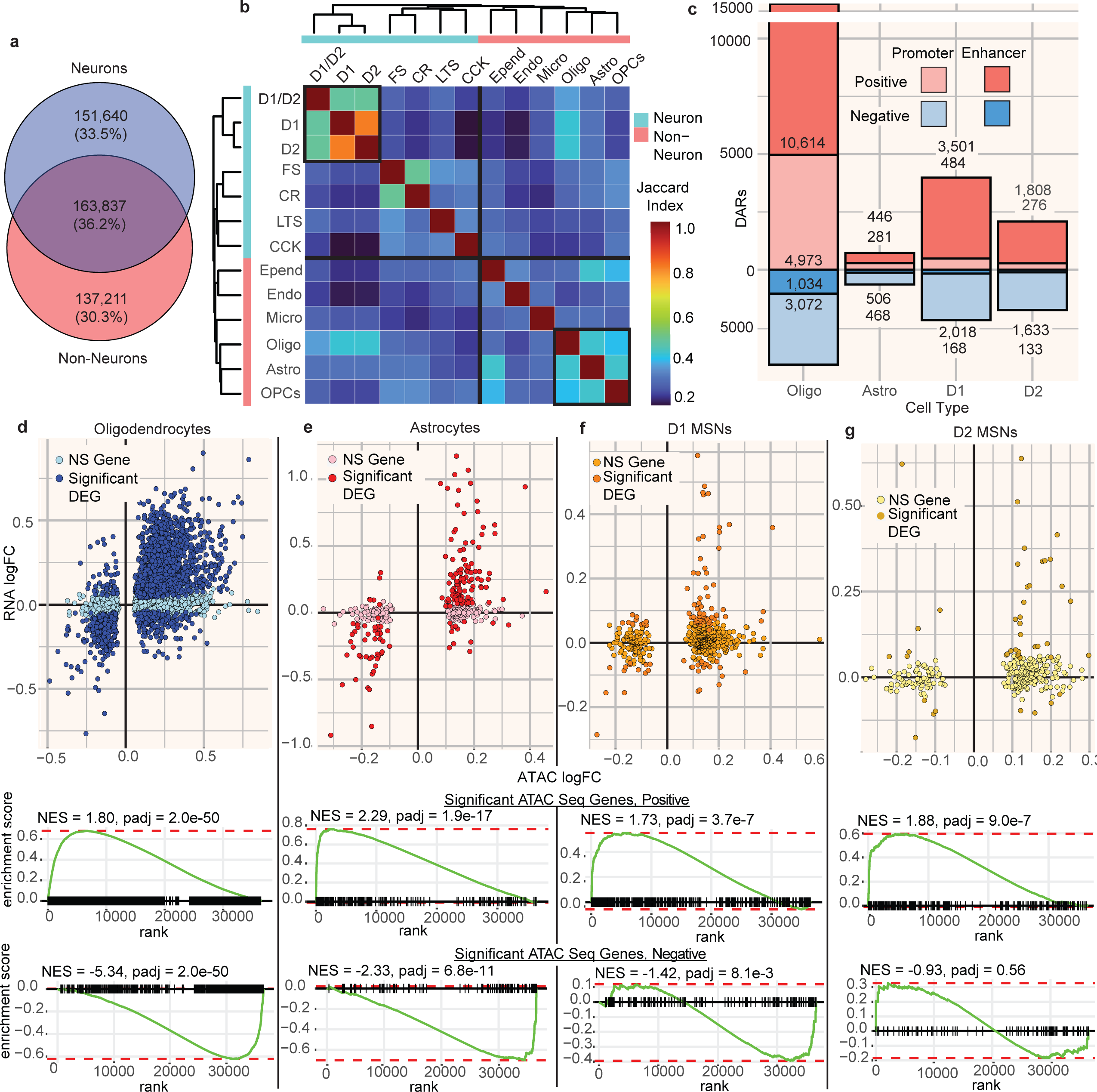
Characterization of AUD-associated changes in chromatin accessibility in the caudate nucleus. **a,** Venn diagram of overlap between the union of open chromatin regions from all neuronal cell types and the union of regions from all non-neuronal cell types. **b,** Heatmap of Jaccard similarity between open chromatin regions in each cell type, hierarchically clustered by Jaccard similarity. **c,** Barplot showing number of differentially accessible regions identified in oligodendrocytes, astrocytes, D1, and D2-type MSNs; red and blue indicate positively and negatively differentially accessible regions, respectively, and lighter and darker coloring indicate regions in promoter and enhancer regions of genes, respectively. Promoter regions were defined as 1 kilobase surrounding the transcription start side of each gene.**d-g,** Top, scatterplot of ATAC peak log(2) fold changes and RNA-seq log(2) fold changes for genes with at least one differentially accessible region (padj < 0.2). Genes are colored based on whether the gene is also differentially expressed (padj < 0.2). Bottom, GSEA enrichment plot of enrichment of the same ATAC-significant genes, split into two sets based on positive or negative effect size, across genes ranked by differential expression fold change. Normalized enrichment scores (NES) and Benjamini-Hochberg adjusted p values (padj) for each GSEA test are shown; d, oligodendrocytes; e, astrocytes; f, D1-type MSNs; g, D2-type MSNs.

To determine the AUD-associated differences in chromatin accessibility for each cell type, we calculated the differentially accessible chromatin regions (DARs) – i.e., open chromatin regions that differed in accessibility between individuals with AUD and those without, again using a pseudobulk method. Samples were removed on a cell type-specific basis if the sample contained less than 50 cells of that cell type (See Supplementary Table 12 for a summary of the number of pseudobulk samples created for each cell type). We identified DARs for eight cell types (Supplementary Table 12); of those, only oligodendrocytes, astrocytes, D1 neurons, and D2 neurons had over 50 DARs (*padj* < 0.2; Fig. 4C). Just as with the DEGs, most of the differences were in the positive direction – chromatin was on average more open in samples from individuals with AUD. However, most of these chromatin accessibility differences were relatively small – only in oligodendrocytes did any DARs surpass an absolute log_2_ fold change of 0.5.

We compared the magnitude and direction of chromatin accessibility differences to the gene expression differences in genes that had at least 1 DAR in the promoter region, a total of 4,915 genes across all cell types. The AUD-associated DARs and DEGs were in the same direction for most genes in the four largest cell clusters (88%, 90%, 73%, and 77% in oligodendrocytes, astrocytes, D1 neurons, and D2 neurons, respectively), and genes containing DARs were enriched among DEGs in the same four cell types (*padj* < 1e-8; Fig. 4D-G). These results together suggest that AUD-associated differences in chromatin accessibility can potentially lead to a corresponding change in *cis*-gene expression.

### Identifying Genes Contributing to AUD Risk

To find genes likely to contribute to AUD risk, we performed an integrative analysis by combining the differentially expressed gene list with GWAS findings and cell type-specific eQTL loci. Our assumption is that if a genetic variant is associated with the expression levels of a nearby gene (eQTL) and the gene locates in a GWAS locus of an AUD-related trait, the increased or decreased expression level of the gene is more likely to contribute to the trait. Several large-scale GWAS have found genetic loci associated with AUD-related traits: 496 independent loci associated with number of drinks per week,^3^ and 90 independent loci associated with PAU^5^, including 5 loci associated with both traits. There are 3,406 and 749 genes, respectively, within these loci, of which 147 were associated with both traits, a total of 4008 unique genes (Supplementary Table 13). Of these, 518 were differentially expressed (*padj* < 0.2) in astrocytes in our snRNA-seq data, 861 in oligodendrocytes, 318 in D1 neurons, and 329 in D2 neurons. Differentially expressed genes within an associated locus are more likely than the others to play a role in these traits. Among these, genes whose expression is also associated with a nearby genetic variant are even more likely to be potential driver genes for AUD (Fig. 5A). Variants within open chromatin near such differentially expressed genes are strong candidates for those that affect the traits.

**Figure 5:**
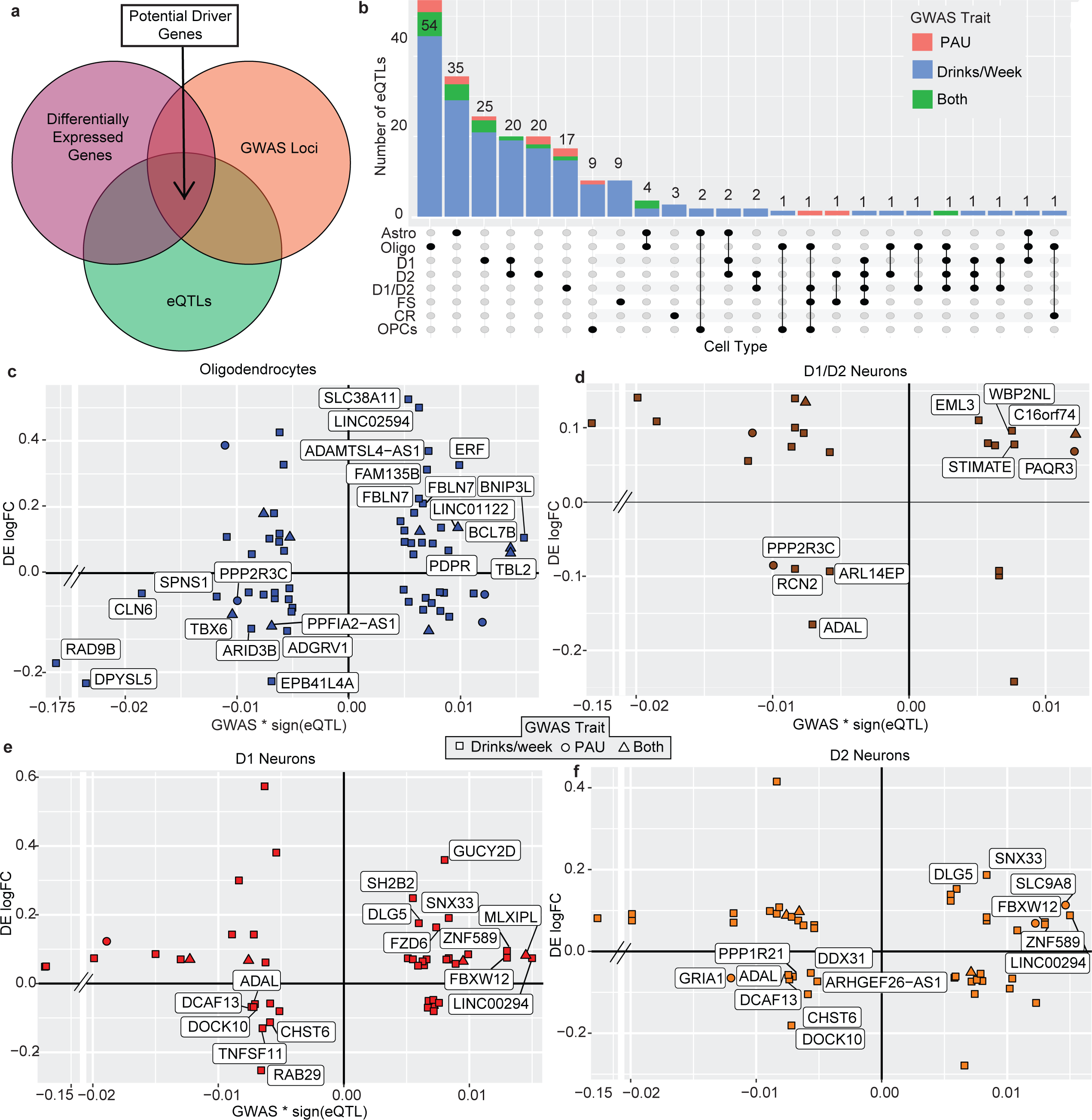
Integration of eQTL analysis with GWAS data and differential gene expression. **a,** Overview of DEG-GWAS-eQTL integration. Potential driver genes in AUD are defined as genes that are differentially expressed, contain GWAS loci associated with an alcohol-related trait, and contain an eQTL. **b,** Upset plot showing number of differentially expressed genes (FDR < 0.2) containing a cis-eQTL (FDR < 0.2) and a GWAS locus for combinations of cell types. The phenotype trait of the GWAS loci overlapping the gene is indicated in color. PAU = Problematic alcohol use. **c,** For oligodendrocytes, differential expression log fold change (individuals with AUD vs those without) plotted against GWAS effect size (using the variant within the locus with the smallest p-value), multiplied by the eQTL effect size, for each gene with a significant eQTL (Benjamini-Hochberg adjusted p-value < 0.2). Shape indicates with which phenotypic trait the GWAS locus overlapping the gene is associated. **d, e, f,** as (c), for oligodendrocytes (d), D1 neurons (e), and D2 neurons (f).

We identified genotypes of the 143 individuals using our snATAC-seq data. For all cell types with at least one DEG, we tested variants overlapping cell type-specific open chromatin regions for an association with expression of nearby differentially expressed genes (see Online Methods). In nine of the eleven cell types tested, we found at least one gene containing a variant in our snATAC-seq data overlapping an open chromatin region that was associated with a differentially expressed GWAS-associated gene. Six cell types contained over 10 genes with such an eQTL (Fig. 5B), and some genes contained eQTLs in multiple cell types. For example, *PPP2R3C,* within a locus associated with PAU, was associated with an eQTL with a negative effect size at variant rs1056879 in oligodendrocytes, D1/D2 MSNs, and fast-spiking interneurons (Extended Data Fig. 7). *PPP2R3C* was expressed at lower levels in both cell types from individuals with AUD. Combined, these findings suggest that AUD is associated with downregulation of this gene’s expression. *ADAL*, in a locus positively associated with alcohol drinks/week, was differentially expressed in D1, D2, D1/D2 and fast-spiking neurons, with a *cis*-eQTL at rs3742971 negatively associated with *ADAL* expression in all four of these cell types (Extended Data Fig. 7).

For a specific variant in the regulatory region, when the direction of the GWAS effect is in the same or opposite direction as its association with the expression of a nearby gene, the higher or lower expression level of the gene may contribute to the trait. These genes are considered risk or protective genes to AUD. We therefore compared the log_2_ fold change of differentially expressed genes with the GWAS effect size multiplied by the sign of the eQTL effect size of the same gene. We identified multiple genes in several cell types that had expression changes in the same direction as the GWAS*eQTL effect, including *PPP2R3C* in oligodendrocytes and D1/D2 MSNs (Fig. 5C-D), and *ADAL* in all MSNs (Fig. 5D-F). The expression of these genes was either positively associated with AUD (genes in the first quadrant) – i.e., risk genes, like *ERF* in oligodendrocytes – or negatively associated (genes in the third quadrant) – i.e., protective genes, such as *PPP2R3C* in oligodendrocytes.

### Cell Type-specific Gene-Regulatory Mechanisms in AUD

To determine which transcription factors and their target genes become more or less active in individuals with AUD, we used LINGER,^43^ a recently developed tool for inferring gene regulatory networks from paired single-cell expression and chromatin accessibility data (see Online Methods). We pooled the pseudobulk data tested for differential accessibility across all cell types to construct the regulatory network. From this network, we extracted key transcription factor-target gene subnetworks (modules). Several regulatory modules were significantly enriched for genes from the two GWAS studies utilized in our analyses. Modules 1, 2, 3, and 10 were enriched for genes associated with PAU,^5^ and modules 3 and 8 were enriched for genes associated with drinks per week^3^ (Fig. 6A).

**Figure 6:**
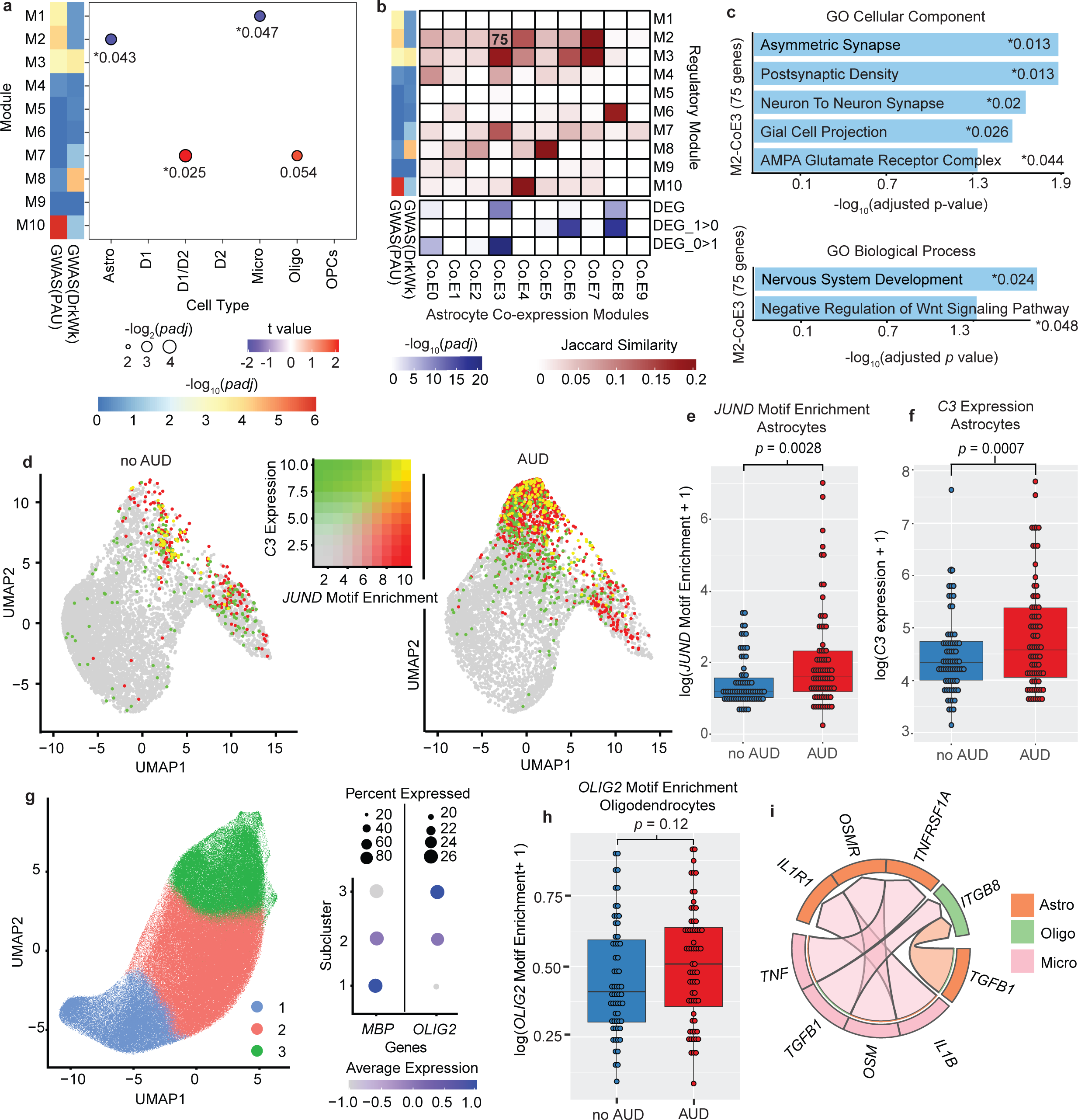
Cell type-specific gene regulatory networks associated with AUD. **a,** Gene regulatory network analysis of samples from individuals with and without AUD comparing the average expression of genes within 10 regulatory modules (M1-M10; see Online Methods). Dot size represents Benjamini-Hochberg adjusted p-value and color represents t-value of the difference in average gene expression. Asterisk indicates significance (<0.05). Left two columns display enrichment of module genes in the set of genes significantly associated with problematic alcohol use (GWAS(PAU)) and drinks per week (GWAS(Drkwk)). **b,** Astrocyte co-expression modules. Left two columns are the same as (a). Top (red heatmap), Jaccard index between genes belonging to the 10 regulatory modules, and 10 co-expression modules (Co.E0-Co.E9), as calculated by WGCNA. Bottom (blue heatmap), Benjamini-Hochberg adjusted p-value of a t test of the difference in average expression of genes in each co-expression module, between individuals with and without AUD. DEG_1>0 indicates higher expression in samples from individuals with AUD, and DEG_0>1 indicates higher expression in those without AUD. Bold number “75” indicates number of genes overlapping between the regulatory module and co-expression module. **c,** Gene Ontology (GO) functional enrichment for the 75 genes overlapping regulatory module 2 and co-expression module 3 shown in (b). Numbers marked by asterisk indicates Benjamini-Hochberg adjusted adjusted p-value of enrichment. **d,** UMAP of astrocytes from individuals without AUD (left plot) and with AUD (right plot). Dot color indicates the enrichment of the *JUND* motif (red) and the log-normalized *C3* expression (green). Yellow indicates high expression of *C3* and high *JUND* motif enrichment. **e,** Boxplot of the log of chromvar motif activity score for the *JUND* motif in astrocytes for samples from individuals with and without AUD. **f,** Boxplot of the log of *C3* expression in astrocytes for samples from individuals with and without AUD. **g,** Left, UMAP of oligodendrocytes, clustered and annotated into three subclusters using graph-based clustering. Right, dotplot of *MBP* and *OLIG2* expression for each of the three oligodendrocyte subclusters. Dot size indicates percentage of cells expressing the gene, and dot color indicates average expression of the gene. **h,** Boxplot of the log of chromvar motif activity score for the *OLIG2* motif in oligodendrocytes for samples from individuals with and without AUD. **i,** Circos plot showing the top five ligand-receptor interactions (determined by scaled ligand activity score from MultiNicheNet) between astrocytes, oligodendrocytes, and microglia.

Among the 10 modules detected, several showed differential expression between those with and without AUD (Fig. 6A, Supplementary Table 14): module 2 genes in astrocytes, module 7 in D1/D2 MSNs (also marginally significant in oligodendrocytes, *padj* = 0.054), and module 1 in microglia (*padj* < 0.05). Module 7 target genes were enriched for several pathways from GP Cellular Component database, including collagen-containing extracellular matrix (*padj* = 0.003), vesicle (*padj* = 0.007) and basement membrane (*padj* = 0.007).

Notably, genes in module 2 in astrocytes and in module 1 in microglia both had lower expression in individuals with AUD and were enriched for genes from the PAU GWAS.^5^ Module 1 contained only one transcription factor, *ZBTB16*, which is a negative regulator of inflammation,^44^ including in microglia.^45,46^ Module 1 contained 6 target genes that were enriched for neurodegeneration in the Human Phenotype Ontology (HP:0002180, *padj* = 0.017), including *KLB*, a gene associated with alcohol consumption.^47^

Module 2, with lower expression in astrocytes from those with AUD, contained 24 transcription factors and 451 target genes. To determine which of these genes were more likely to be regulated together, we performed weighted gene co-expression network analysis (WGCNA^48^) of astrocyte single-nucleus data, which revealed ten groups of co-expressed genes (Supplementary Table 15). Co-expression group 3 (Co.E3) was enriched for differentially expressed genes, particularly those with lower expression in individuals with AUD. There were 75 genes in Co.E3 that overlapped with regulatory module 2 (Fig. 6B), suggesting that these genes not only show similar expression patterns but also have similar patterns of regulation. Functional enrichment analysis identified several enriched pathways, including pathways relating to glutamatergic synapses, nervous system development, and negative regulation of Wnt signaling pathways from GO Biological Process and Cellular Component (Fig. 6C).

Next, we identified several key regulatory transcription factors in each cell type using the expression and accessibility of all target genes within the trans-regulatory network constructed by LINGER^43^ (Extended Data Fig. 8). For example, in D2 medium spiny neurons, the transcription factor predicted to have the most significant altered activity in AUD, based on the chromatin openness of its target genes, was *FOXO1* (Forkhead transcriptional factor O1, Extended Data Fig. 8), which has been previously shown to regulate energy homeostasis in neurons^49^ and was linked to depression in multiple recent studies.^50,51^

In astrocytes, six transcription factors were implicated in regulating AUD-associated differences in both expression and chromatin accessibility, including the gene *JUND* (JunD proto-oncogene), which had increased regulatory activity in individuals with AUD in astrocytes (Extended Data Fig. 8). We found 142 transcription factor motifs with significantly higher enrichment in those with AUD (*p* < 0.05) using chromVAR.^52^ 15 of the top 20 enriched motifs (by p-value) with significantly higher activity in cells from individuals with AUD are annotated by JASPAR^53^ as being associated with bZIP transcription factors (*p* <= 0.005; Supplementary Table 16). The bZIP transcription factor motif for JUND had significantly higher enrichment in astrocytes in individuals with AUD (*p* = 0.01, Fig. 6D-E). Astrocytes with high *JUND* motif enrichment were largely in regions with high complement component 3 (*C3*) expression, a marker of reactive astrogliosis.^54^ There was higher expression of *C3* in individuals with AUD (1.65 fold, *padj* = 0.0007; Fig. 6F).

We saw a modest decrease in expression of myelin basic protein (*MBP*), a major component of myelin, in oligodendrocytes from individuals with AUD (*padj* = 0.11). Graph-based subclustering of oligodendrocytes revealed three major subclusters, one of which had significantly lower *MBP* expression than the other two (Fig. 6G). This cluster was marked by high expression of *OLIG2*. We found 616 motifs with activity significantly associated with AUD (*p* < 0.05), including *OLIG2* (*p =* 0.028, Supplementary Table 17) (Fig. 6H). This gene was identified as a ‘driver’ by LINGER, having higher regulatory activity in individuals with AUD based on the chromatin accessibility of its target genes.

Several genes implicated in the gene regulatory analysis were also implicated in our integrative GWAS/eQTL/differential expression analyses, including three target genes in astrocytes from regulatory module 2: *BTBD3*, *LRRC4C*, and *PTBP2* (Extended Data Fig. 9). Because these three genes are (1) differentially expressed in AUD, (2) part of a differentially expressed regulatory module identified in gene regulatory network analysis, (3) *cis* to a variant associated with gene expression, and (4) associated with drinks per week, they have strong evidence linking them as potential driver genes of alcohol consumption and/or AUD in the caudate.

### Activated Microglia Induce Reactive Astrocytes and Oligodendrocytes

We measured changes in ligand-receptor signaling pathways between microglia, astrocytes, and oligodendrocyte cells using MultiNicheNet^55^. We identified three ligand-receptor pairs that signal from microglia to astrocytes with high downstream gene activity: *IL1B-IL1R1*, *OSM*-*OSMR*, and *TNF*-*TNFRSF1A* (Fig. 6I, see Supplementary Table 18 for all ligand-receptor pairs). These three pairs have been shown to work synergistically to induce pro-inflammatory cytokines in astrocytes.^56^ Predicted downstream target genes of *IL1B*-*IL1R1* in astrocytes included *C3*, the widely used marker of reactive astrocytes, as well as bZIP family TFs *FOSL1*, *XBP1*, and *CEBPD.* We identified a single ligand-receptor pair from both astrocytes and microglia to oligodendrocytes with a high ligand activity: *TGFB1*-*ITGB8* (Extended Data Fig. 10).

## Discussion

In this study we present the first integrated profile at the single-nuclei level of differences between individuals with and without AUD in gene expression, chromatin accessibility, and cell state in the caudate nucleus, and determined potential regulatory mechanisms underlying these differences. By combining single-nucleus RNA sequencing (RNA-seq) with chromatin accessibility profiling (ATAC-seq) within the same cells at large scale, using both sn-multiome and snRNA-seq experiments, we discovered novel patterns of gene expression within different cell types and novel regulatory mechanisms behind them. This integrative dual approach demonstrates the power of large-scale multiomic studies to uncover cell-type-specific regulatory mechanisms in complex brain disorders. Such methodologies can be applied to investigate other neuropsychiatric and neurodegenerative conditions, expanding the utility of this approach across the field of neuroscience.

Our large sample size allowed us to identify 17 distinct cell types in the human caudate, including rare interneuron and non-neuronal populations not previously identified in single-cell studies such as a recent single-cell atlas of the brain^57^. These include a population of cholecystokinin/vasoactive intestinal polypeptide-expressing neurons detected in small numbers in animal models but previously not detected in the human striatum,^33^ a population of calretinin-expressing neurons, knowledge of which has been extremely limited,^33^ and a small cluster of vascular smooth muscle cells, a cell type which has recently been linked to neurovascular coupling;^58^ a recent study linked neurovascular coupling with chronic alcohol exposure in mice.^59^

We identified thousands of genes differentially expressed in a range of cell types and characterized the accompanying differences in chromatin accessibility. The differentially expressed genes and differentially accessible regions were highly correlated between D1- and D2-type medium spiny neurons, which are components of the direct and indirect pathways, respectively, of the basal ganglia. Pathways enriched in individuals with AUD related to RNA processing (‘RNA Metabolism’, ‘Processing of Capped Intron-Containing pre-mRNA’) and immune response (‘Innate Immune System’, complement-related pathways). Recent studies have also found dysregulation of neuroimmune genes in neurons in several brain regions in mice.^60^, Gene regulatory network analysis allowed us to identify transcription factors possibly regulating these differences. For example, *FOXO1* was shown to have significantly lower activity in D2 medium spiny neurons from individuals with AUD, based on the chromatin openness of its target genes. *FoxO1* in mice has been linked to regulation of energy homeostasis in neurons,^49^ and a lack of *FoxO1* in the brain caused a depressive-like phenotype.^50,51^ This could represent a novel regulatory gene in neurons in AUD, as AUD and major depression frequently co-occur.^61^

We observed a small cluster of medium spiny neurons expressing both D1 and D2 dopamine receptor genes. This MSN type, variously described as eccentric MSNs, D1H, and D1/D2 hybrid MSNs,^41,62^ has been observed in mice,^62,63^ primates,^41^ and humans,^64^ but its association with AUD has thus far been unknown. The pattern of gene expression changes in these neurons was less correlated with the other MSN types and different biological pathways were enriched: ‘HDMs Demethylate Histones’ was the most highly enriched pathway in individuals with AUD, while many metabolic and translational control pathways had lower than expected expression in those with AUD. These differences from the classical D1 and D2 neurons suggest that D1/D2 hybrid MSNs play a distinct role in the caudate. These neurons have been shown in a recent study in mice to be morphologically distinct from D1 and D2-type MSNs, with a smaller cell body, less expansive dendrite structure, and fewer spines, and were differently affected by treatment with a denervating agent.^63^ Genes involved in collagen-containing extracellular matrix and vesicular pathways were upregulated in those with AUD. Collagen is a major component of the blood-brain barrier,^65^ so these neurons could play a role in its integrity or regulation, processes which may be disrupted in AUD.

We identified four distinct states of microglial cells that did not adhere closely to the classical M1/M2 (inflammatory/anti-inflammatory) distinction^66^, although some of these cell clusters were enriched for markers of inflammation Indeed, recent studies are beginning to question the biological accuracy of the widely-cited M1/M2 classification.^67^ One subcluster was distinguished by high *CD83* expression, a gene implicated in regulating both inflammatory and anti-inflammatory processes.^39^ An increased proportion of microglia showed an inflammatory gene expression profile in those with AUD. Chronic alcohol exposure has been shown to cause microglial activation in mice, leading to neuroinflammation.^68^ Our analyses found that expression of *ZBTB16,* a negative regulator of inflammation,^44^ was a key regulator of gene module 1, and *ZBTB16* expression was decreased in those with AUD. Thus, the increased inflammatory response in microglia in AUD could be due in part to decreased *ZBTB16* activity. *ZBTB16* is known to counteract microglial M1 activation,^45^ and its knockout in mice caused increased microglia and autism-like and schizophrenia-like behaviors,^46^ but it had not been implicated in AUD.

In astrocytes from individuals with AUD there was significantly higher expression of astrocyte reactivity marker *C3*. This extends to the caudate evidence from studies that demonstrated that inflammation evoked by ethanol exposure is accompanied by reactive astrogliosis.^8,69^ The cells with increased *C3* expression also had significantly higher predicted activity of bZIP transcription factors such as *JUND,* suggesting that changes in these transcription factors may be regulating astrocytes as they undergo reactive astrogliosis. Gene regulatory network analysis identified a group of genes with similar patterns of *trans-*regulation that had decreased expression in AUD. These genes were overrepresented within gene sets related to glutamatergic synapses, consistent with previous work linking the disruption of glutamate homeostasis in astrocytes to AUD.^69^ Other enriched pathways in these co-expressed and regulatory genes in astrocytes include nervous system development and negative regulation of Wnt signaling. A recent study implicated the disruption of Wnt signaling in the striatum of rats as an effect of high cocaine self-administration,^70^ but its potential link with alcohol is novel. Together this suggests that decreased regulation of Wnt signaling, especially in astrocytes, may play an important role in alcohol addiction.

In oligodendrocytes, we observed differences in expression and chromatin accessibility in thousands of genes, and enrichment in biological pathways relating to neurotransmitter uptake and depolarization. The gene encoding myelin basic protein had slightly, but significantly, lower expression in individuals with AUD. There is evidence that action potential propagation through axons can be regulated by oligodendrocyte depolarization.^71^ In pathological conditions such as excitotoxicity, excessive neurotransmitter release from neurons can lead to an excessive intracellular Ca^2+^ flux into oligodendrocytes, damaging myelinating processes.^72^ Lower *MBP* expression was limited to cells marked by higher expression of *OLIG2* (Fig. 6G-H). *OLIG2*, a master regulator in mature and developing oligodendrocytes, has been shown to have higher activity after brain injury,^73^ and is linked to myelination: replacing *Olig2* with its dominant-active form in rodents led to decreased expression of *MBP,*^74^ and deletion of the *Olig2* gene accelerated remyelinating processes.^75^ This suggests that our observed increase in *OLIG2* activity in individuals with AUD may in part lead to dysregulation of myelination in oligodendrocytes. Indeed, alcohol consumption and alcohol use disorder have been found to be associated with white matter degeneration,^76,77^ but prior to this study, there had not been a direct link between AUD, demyelination, and specific genes such as *OLIG2*.

The AUD-associated chromatin differences were correlated with expression differences, as expected. In every cell type, genes implicated in immune pathways – such as cytokine/interferon response, innate immune system, and complement cascades – were overrepresented among those differentially expressed. This extends previous findings that chronic alcohol exposure is associated with an increased neuroimmune response in neurons, glia and astrocytes.^78,24^

Leveraging single-cell transcriptomic data to identify AUD-associated differences in communication pathways between cell types allowed us to integrate the transcriptomic and epigenetic alterations in each cell type into predictions of upstream signaling events. Signaling from microglial cells to astrocytes that involves proinflammatory molecules IL-1β, TNF, and oncostatin M is higher in individuals with AUD, concordant with the hypothesis that activated microglia induce neurotoxic reactive astrocytes.^54^

These three molecules have been shown to work synergistically in astrocytes and other cells to induce pro-inflammatory and neurotoxic molecules such as nitric oxide^79^ and prostaglandin E(2).^56^ Although reactive astrocytes can induce death of neurons and oligodendrocytes, we did not observe a significant difference between individuals with and without AUD in relative proportions of neuronal cell types or oligodendrocytes. We found increased signaling via *TFGB1-ITGB8* from both microglia and astrocytes to oligodendrocytes. TGF-*β*1 signaling is known to increase after injury, and studies have shown that ethanol exposure induces TGF-*β*1 signaling in rats.^80,81^ Previous work showed that TGF-*β*1 expression increases in astrocytes and microglia in animal models of cerebral ischemia,^82^ and another study has shown that TGF-β1 signaling plays a role in myelination in oligodendrocytes.^83^

The nature of AUD does not normally allow us to differentiate between pre-existing genetic differences and those due to the chronic alcohol consumption that is the hallmark of AUD. Some AUD-associated differences we observed (e.g., changes in inflammatory and myelinating processes) appear to be associated with the consequences of AUD. However, pre-existing genetic information (GWAS data and eQTL) allows us to identify possible causes of the disease and genes driving the effects we observed and inferred. We found several genes in multiple cell types with strong evidence of being linked to AUD. For example, the variant rs1412825, located within a locus positively associated with drinks per week,^3^ was negatively associated with expression of the gene Serine/threonine-protein phosphatase 2A regulatory subunit B’’ subunit gamma (*PPP2R3C*) in both oligodendrocytes and D1/D2 MSNs. This, combined with our finding that *PPP2R3C* had significantly lower expression in oligodendrocytes and D1/D2 MSNs in individuals with AUD, suggests that *PPP2R3C* could be protective against AUD. Interestingly, expression of *PPP2R3C* was recently shown to be significantly associated with PAU in the nucleus accumbens – another part of the striatum – using a transcriptome-wide association analysis.^15^ A single variant, rs3742971, within a locus positively associated with PAU,^5^ was negatively associated with expression of the gene encoding the adenosine deaminase-like protein (*ADAL*) in four neuronal cell types (D1 MSNs, D2 MSNs, D1/D2 MSNs, and FS interneurons). *ADAL* had lower expression in individuals with AUD in these cells. In mice, elevated *Adal* levels contribute to low alcohol preference.^84^ This suggests that *ADAL* may be an important factor in the development of AUD. Our integrative analyses found three genes in astrocytes with strong evidence of being drivers of AUD. One of these, BTB/POZ domain-containing 3 (*BTBD3*, a transcription factor), has been shown to regulate compulsive-like behavior in mice,^85^ further evidence of this gene’s potential importance in addiction.

There are several limitations to our study. The use of postmortem brain samples means that both pre-existing differences that increase risk for AUD and differences associated with the extended, excessive alcohol consumption characteristic of it are both present. We cannot clearly delineate which differences are consequences and which are causes of AUD, although our incorporation of GWAS and eQTL information allowed us to infer some of the latter. Experimental studies such as high-throughput CRISPR inhibition or activation of the genes identified herein in cell models could confirm some of the networks and regulatory pathways,^86^ but cellular studies will not allow confirmation of effects on behavior of individuals. Another limitation is that those who drink heavily are more likely to smoke. A recent study found that 63.3% of drinkers at risk of alcohol dependence were smokers compared with 18.2% among drinkers not at risk, and 19.2% among non-drinkers.^87^ Thus, some differences might be attributed in part to smoking. Finally, our results are primarily from males genetically closely related to the 1000 Genomes European samples, and therefore do not capture the transcriptional and epigenetic diversity across ancestry groups or sexes.^88^

In conclusion, we provide a detailed picture of the vast transcriptional and epigenetic differences between individuals with and without AUD in many different cell types within the caudate nucleus that illuminates biological mechanisms underlying these differences and identifies potential driver genes causing these differences. Our work adds novel insights into the etiology associated with AUD, pointing to key pathways and regulatory genes, and underscores the potential of large-scale multiomic datasets to provide novel insights into brain regions and diseases.

## Acknowledgements

The Collaborative Study on the Genetics of Alcoholism (COGA), Principal Investigators B. Porjesz, V. Hesselbrock, T. Foroud; Scientific Director, A. Agrawal; Translational Director, D. Dick, includes ten different centers: University of Connecticut (V. Hesselbrock); Indiana University (H.J. Edenberg, T. Foroud, Y. Liu, M.H. Plawecki); University of Iowa Carver College of Medicine (S. Kuperman, J. Kramer); SUNY Downstate Health Sciences University (B. Porjesz, J. Meyers, C. Kamarajan, A. Pandey); Washington University in St. Louis (L. Bierut, J. Rice, K. Bucholz, A. Agrawal); University of California at San Diego (M. Schuckit); Rutgers University (J. Tischfield, D. Dick, R. Hart, J. Salvatore); The Children’s Hospital of Philadelphia, University of Pennsylvania (L. Almasy); Icahn School of Medicine at Mount Sinai (A. Goate, P. Slesinger); and Howard University (D. Scott). Other COGA collaborators include: L. Bauer (University of Connecticut); J. Nurnberger Jr., L. Wetherill, X., Xuei, D. Lai, S. O’Connor, (Indiana University); G. Chan (University of Iowa; University of Connecticut); D.B. Chorlian, J. Zhang, P. Barr, S. Kinreich, G. Pandey (SUNY Downstate); N. Mullins (Icahn School of Medicine at Mount Sinai); A. Anokhin, S. Hartz, E. Johnson, V. McCutcheon, S. Saccone (Washington University); J. Moore, F. Aliev, Z. Pang, S. Kuo (Rutgers University); A. Merikangas (The Children’s Hospital of Philadelphia and University of Pennsylvania); H. Chin and A. Parsian are the NIAAA Staff Collaborators. We continue to be inspired by our memories of Henri Begleiter and Theodore Reich, founding PI and Co-PI of COGA, and also owe a debt of gratitude to other past organizers of COGA, including Ting-Kai Li, P. Michael Conneally, Raymond Crowe, and Wendy Reich, for their critical contributions. This national collaborative study is supported by NIH Grant U10AA008401 from the National Institute on Alcohol Abuse and Alcoholism (NIAAA) and the National Institute on Drug Abuse (NIDA).

Tissues were received from the New South Wales Brain Tissue Resource Centre at the University of Sydney which is supported by the University of Sydney and by the National Institute of Alcohol Abuse and Alcoholism (R28AA012725). Research reported in this publication was supported by the National Institutes of Health under Award Number U10AA008401. The content is solely the responsibility of the authors and does not represent the official views of the National Institutes of Health.

## Author Contributions

Y.L. and H.J.E jointly conceived the study; X.C. and P.M. performed experiments, supervised by X.X., Y.W., and H.G.; N.G. analyzed data and wrote the manuscript, supervised by Y.L., H.J.E., H.G., and J.L.R.; Q.Y. and Z.D. developed and carried out the gene regulatory network analysis. D.L. performed the genotype imputation analysis; G.J. and H.G. performed initial processing of multiome data and called variants from the ATAC-seq data; J.S and G.T.S. provided postmortem samples; All authors contributed to the interpretation and editing of the manuscript.

## Competing Interests

The authors declare no competing interests.

## Materials & Correspondence

Correspondence and material requests should be addressed to Yunlong Liu and Howard Edenberg.

---

**Extended Data Figure 1:**
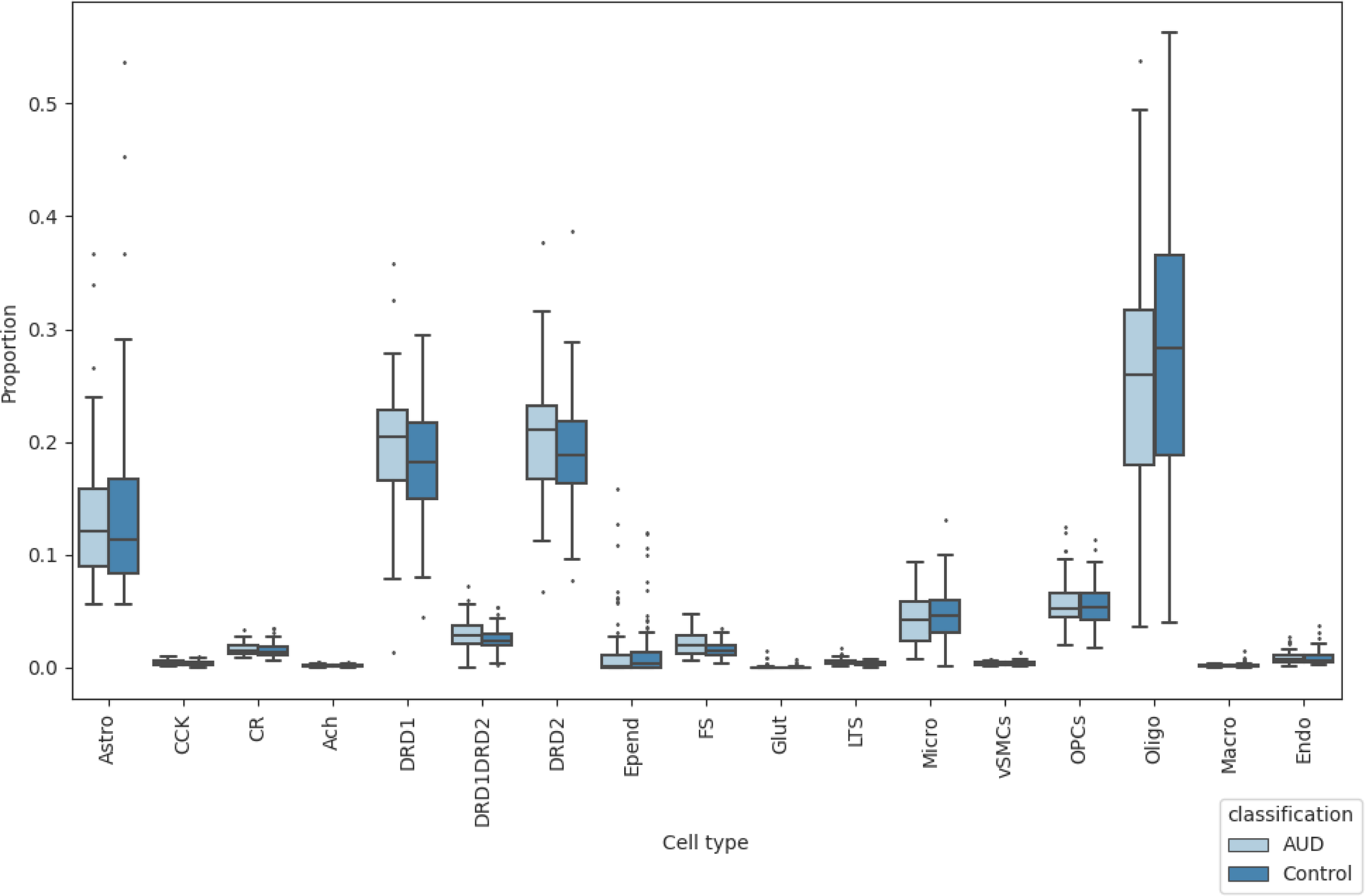
**Proportion of each cell type for each of the 163 samples**, grouped by AUD classification, for all snRNA-seq barcodes used in cell clustering and cell type annotation (see Fig. 1).

**Extended Data Figure 2:**
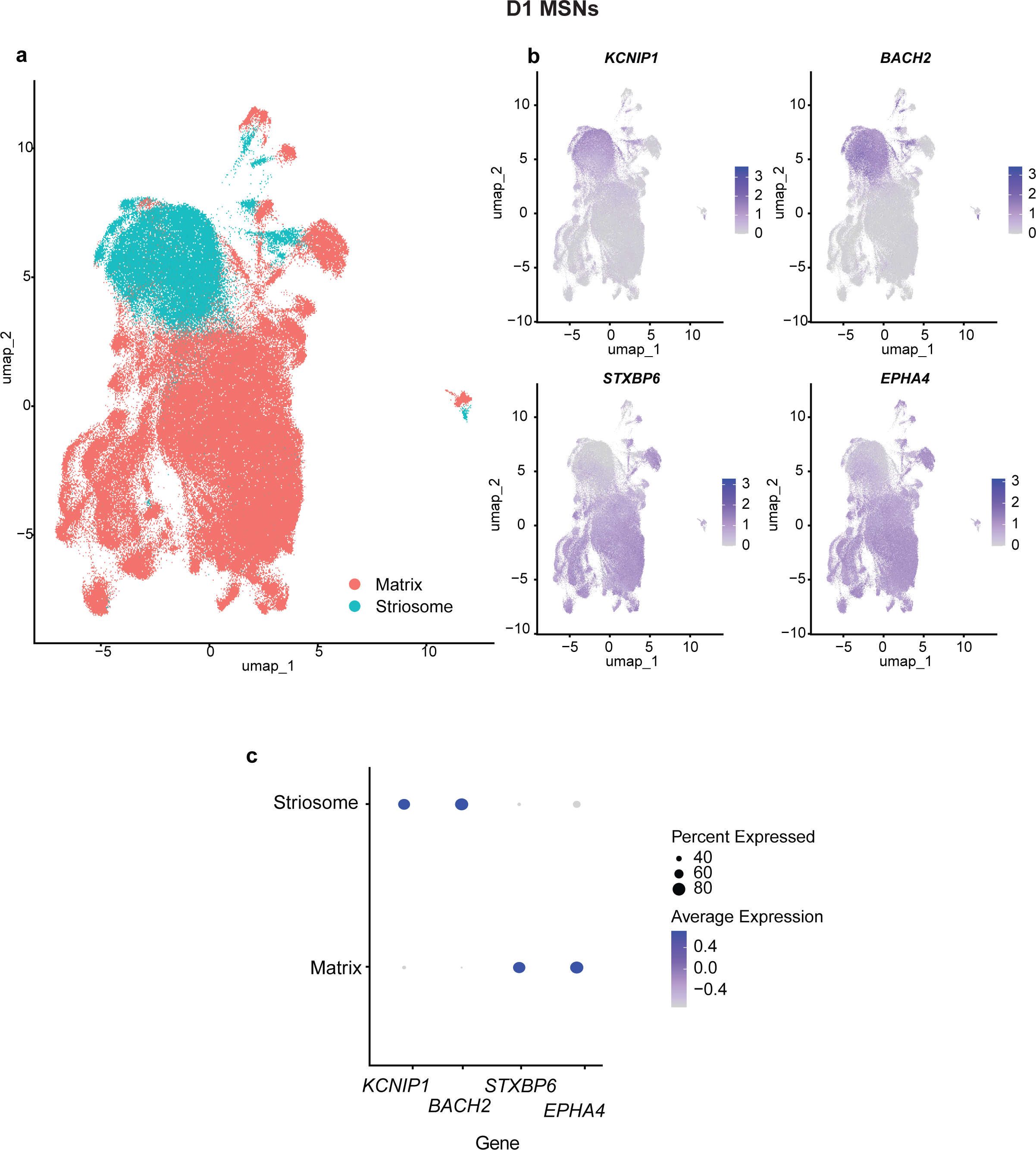
D1 medium spiny neurons subtypes. a, UMAP of D1 MSN cells, colored by compartment (either matrix or striosome). b, UMAP of D1 MSN cells, colored by expression of marker genes used to assign compartment. c, Dotpot of expression and prevalence of representative marker genes for matrix and striosome compartments.

**Extended Data Figure 3.**
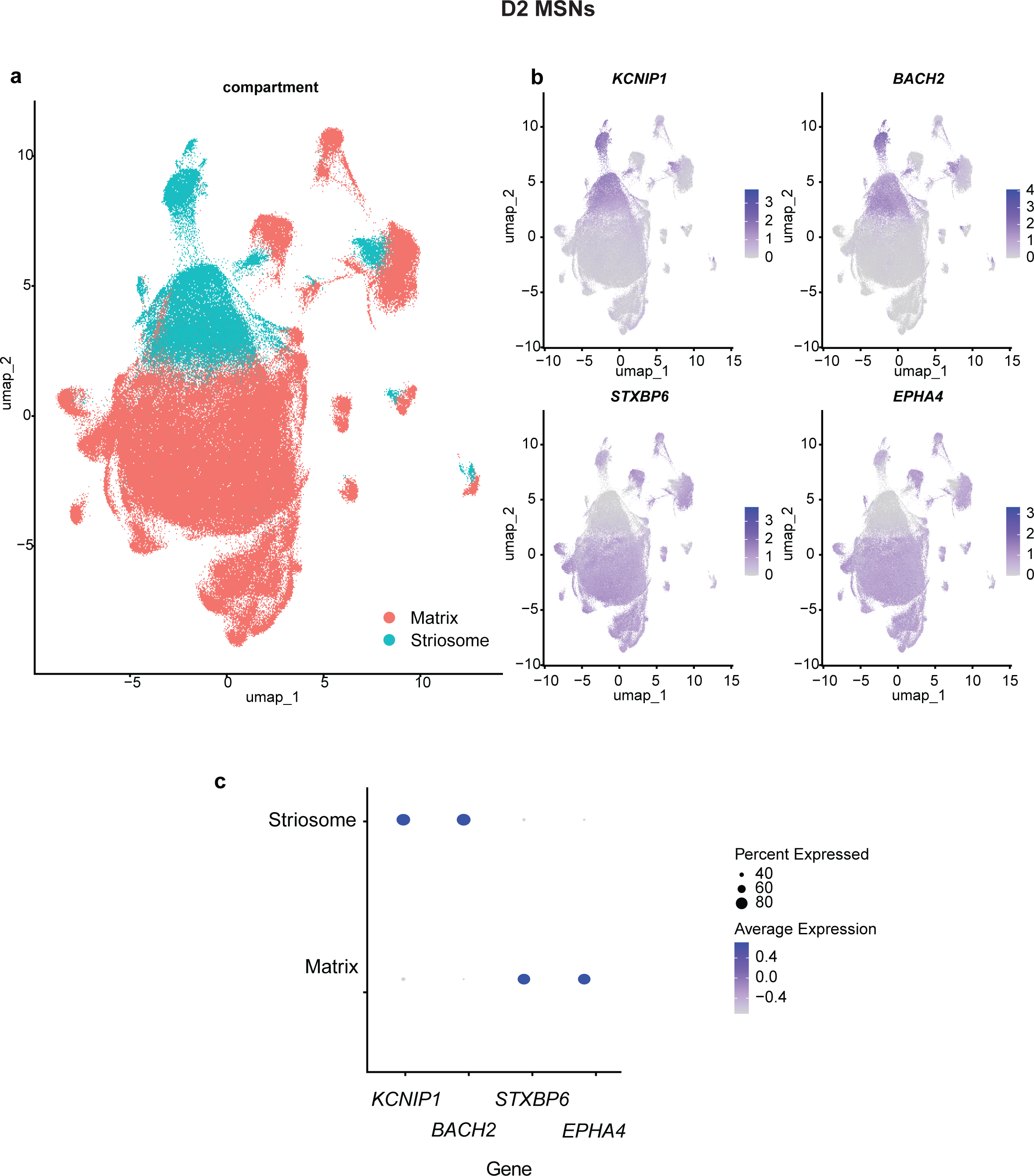
D2 medium spiny neurons subtypes. a, UMAP of D2 MSN cells, colored by compartment (either matrix or striosome). b, UMAP of D2 MSN cells, colored by expression of marker genes used to assign compartment. c, Dotpot of expression and prevalence of representative marker genes for matrix and striosome compartments.

**Extended Data Figure 4:**
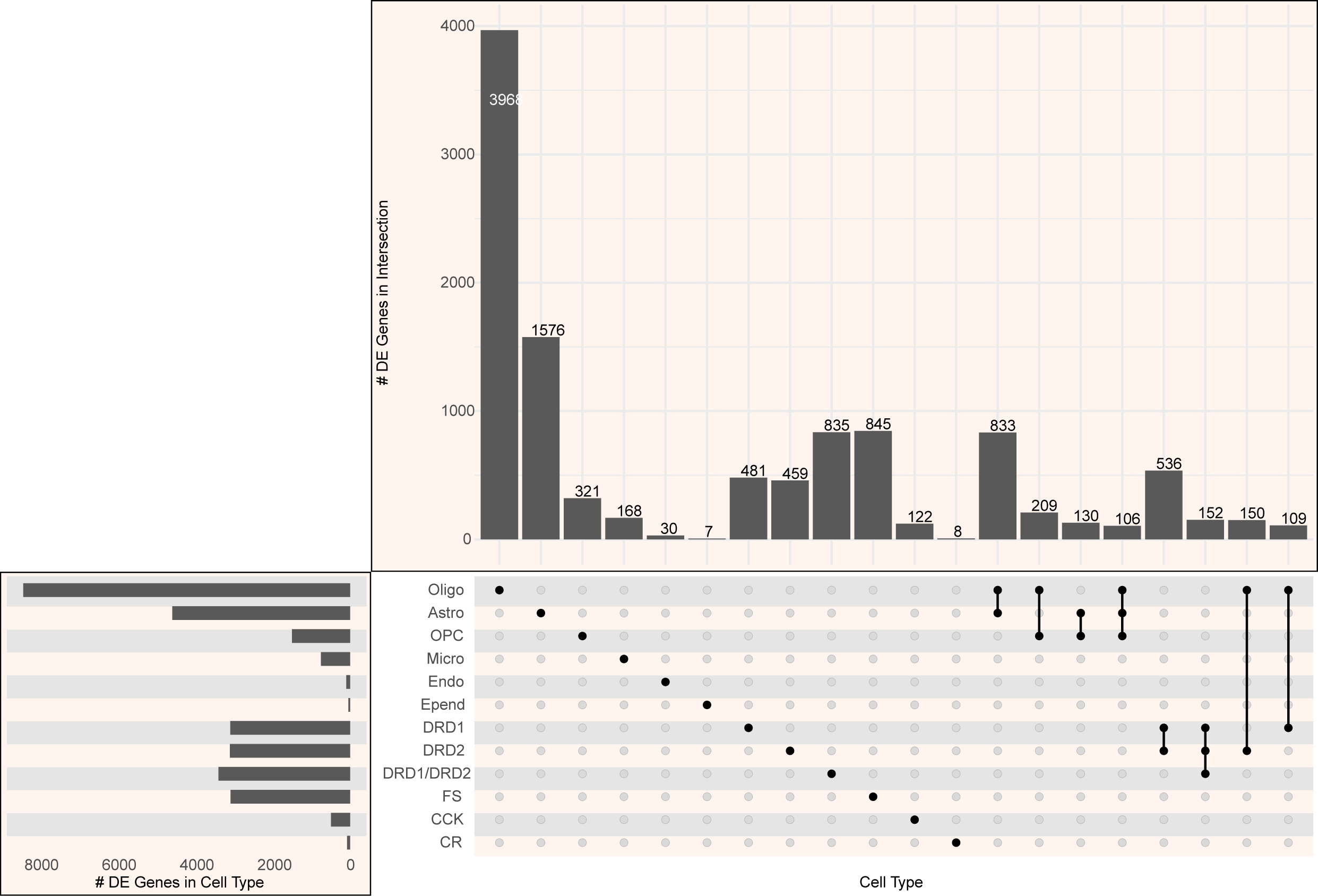
Number of differentially expressed genes in multiple cell types. Upset plot shows the number of genes with expression significantly associated with AUD (padj < 0.2) in different combinations of cell types.

**Extended Data Figure 5:**
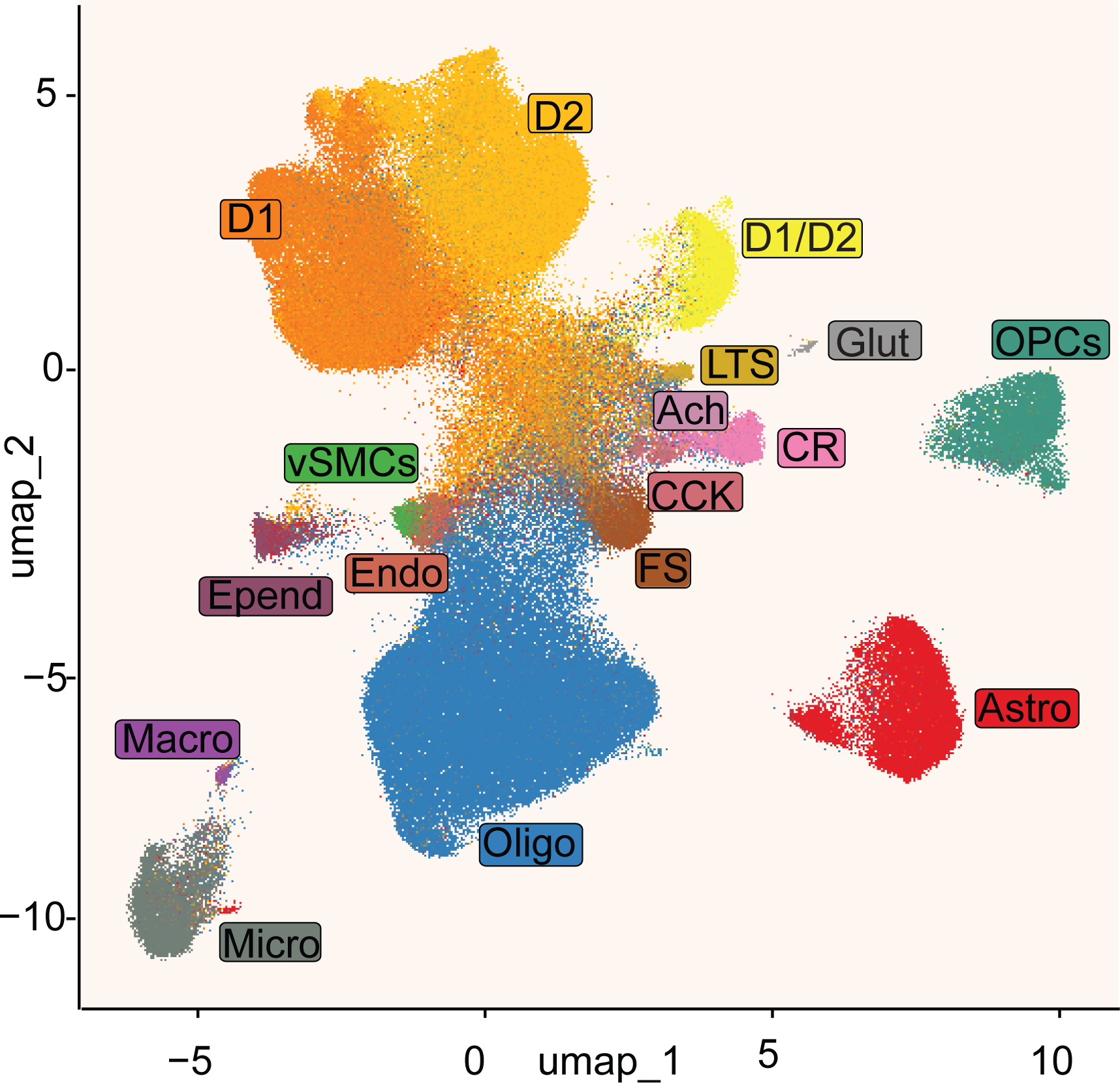
snATAC-seq cell landscape. UMAP plotting each cell for which snATAC-seq data was available, clustered based on snATAC-seq data. Cell type labels for each cell was provided based on the cell’s snRNA-seq data (see Fig. 2).

**Extended Data Figure 6:**
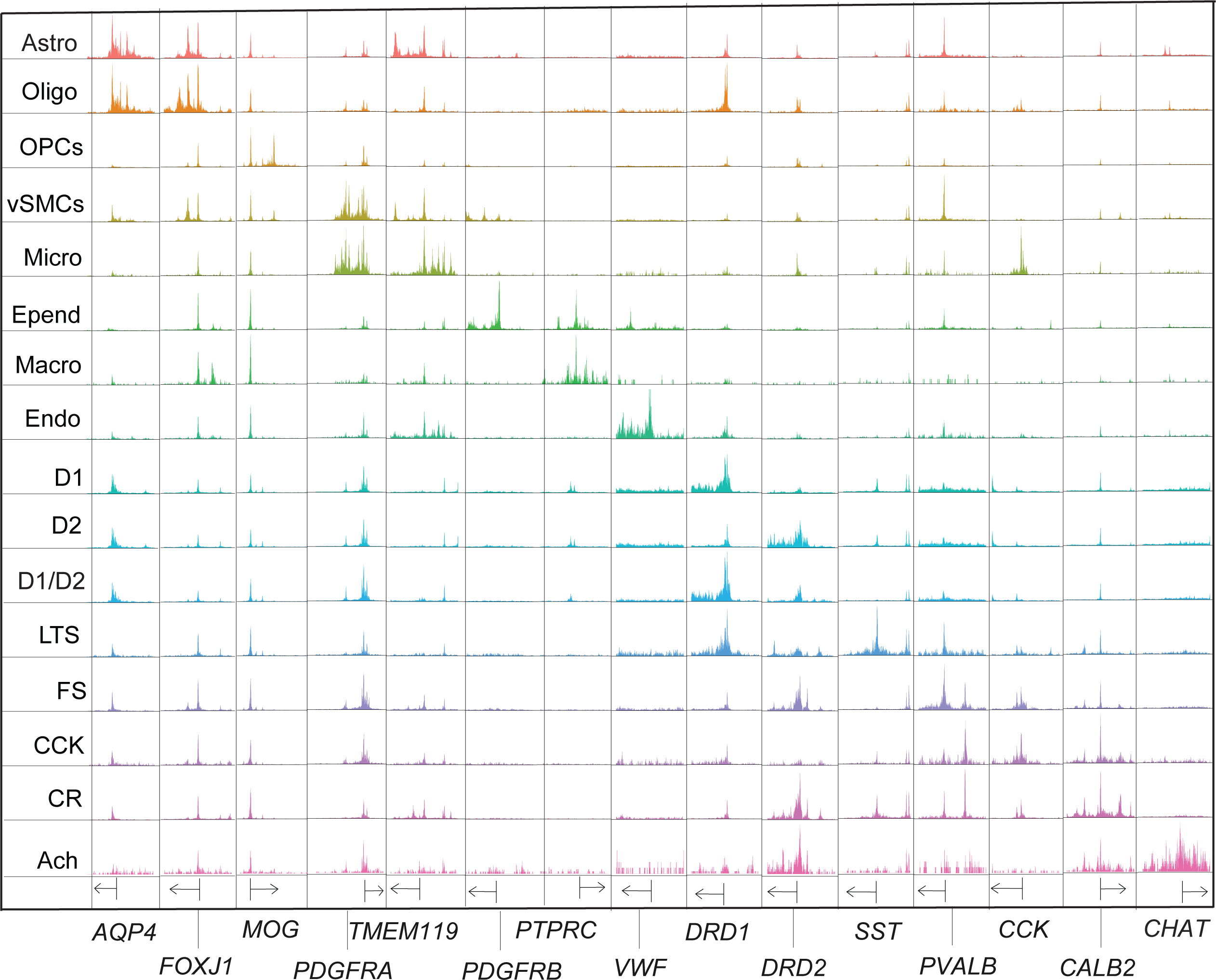
Pseudobulk accessibility profiles for each cell type at canonical marker genes. 5 kilobases on each side of the transcription start site of each gene are shown. Arrow indicates the direction of transcription.

**Extended Data Figure 7:**
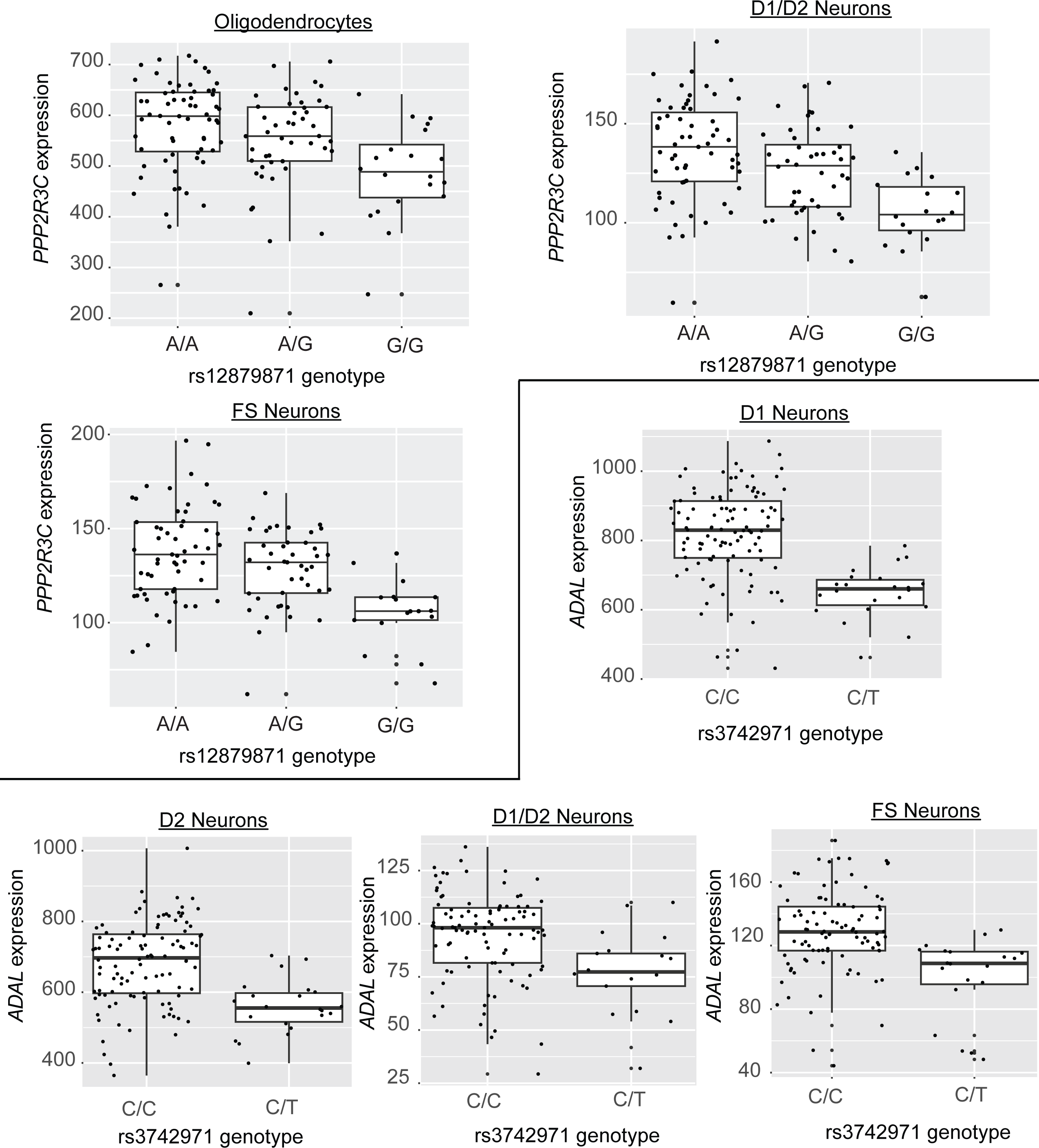
Normalized pseudobulk expression of *PPP2R3C* (above) and *ADAL* (below) for each sample. Plotted by genotype for the significant expression quantitative trait loci (eQTL) in the cell types in which the eQTL was significant.

**Extended Data Figure 8:**
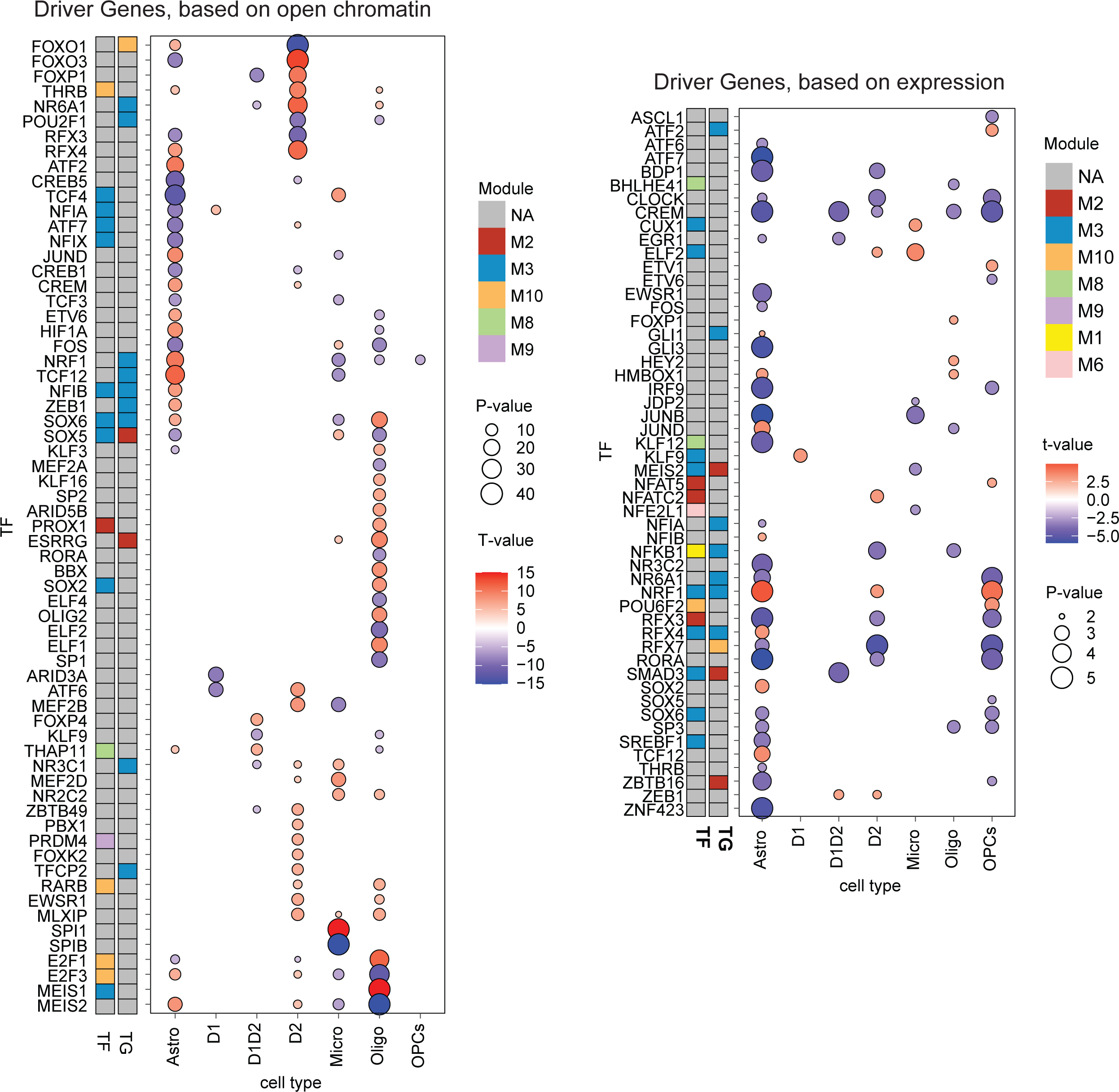
Cell type-specific driver gene scores. **a,** Dotplot of all driver genes found, using LINGER’s driver score, based on chromatin accessibility of target genes of that transcription factor. Size of dot corresponds to p-value and color indicates t-value, of change in driver score between AUD and control individuals. Left two columns correspond to membership in regulatory modules. Genes ordered by cell type with most significant difference in driver score. **b,** Dotplot of all driver genes found, using LINGER’s driver score, based on gene expression of target genes of that transcription factor. Size of dot corresponds to p-value and color indicates t-value, of change in driver score between AUD and control individuals. Left two columns correspond to membership in regulatory modules. Genes are ordered alphabetically.

**Extended Data Figure 9:**
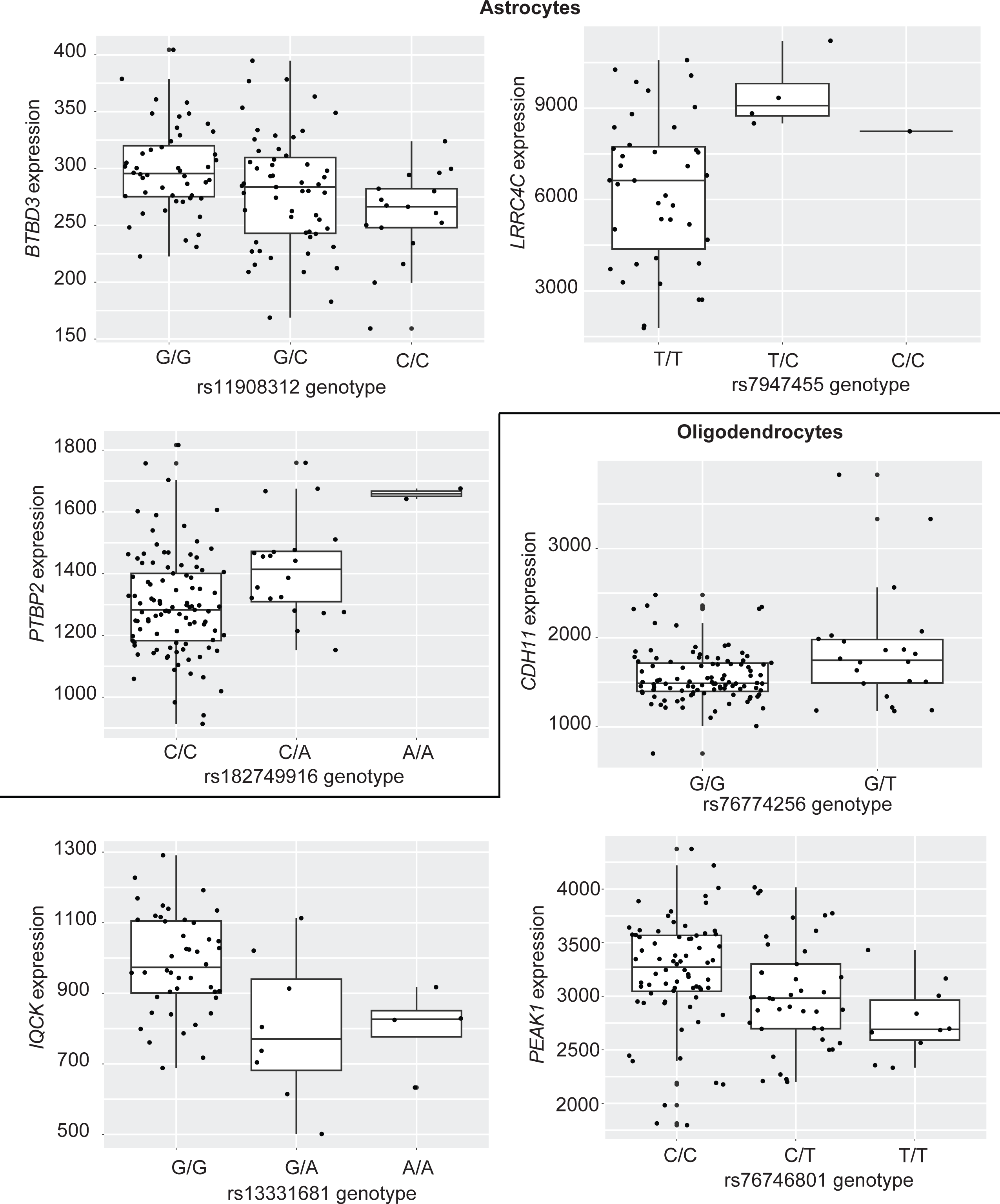
Normalized pseudobulk expression of target genes with eQTLs. Genes from modules 2 and 3 (as identified by LINGER) with a significant eQTL in astrocytes and oligodendrocytes. Plotted by genotype for the signficant eQTLs.

**Extended Data Figure 10:**
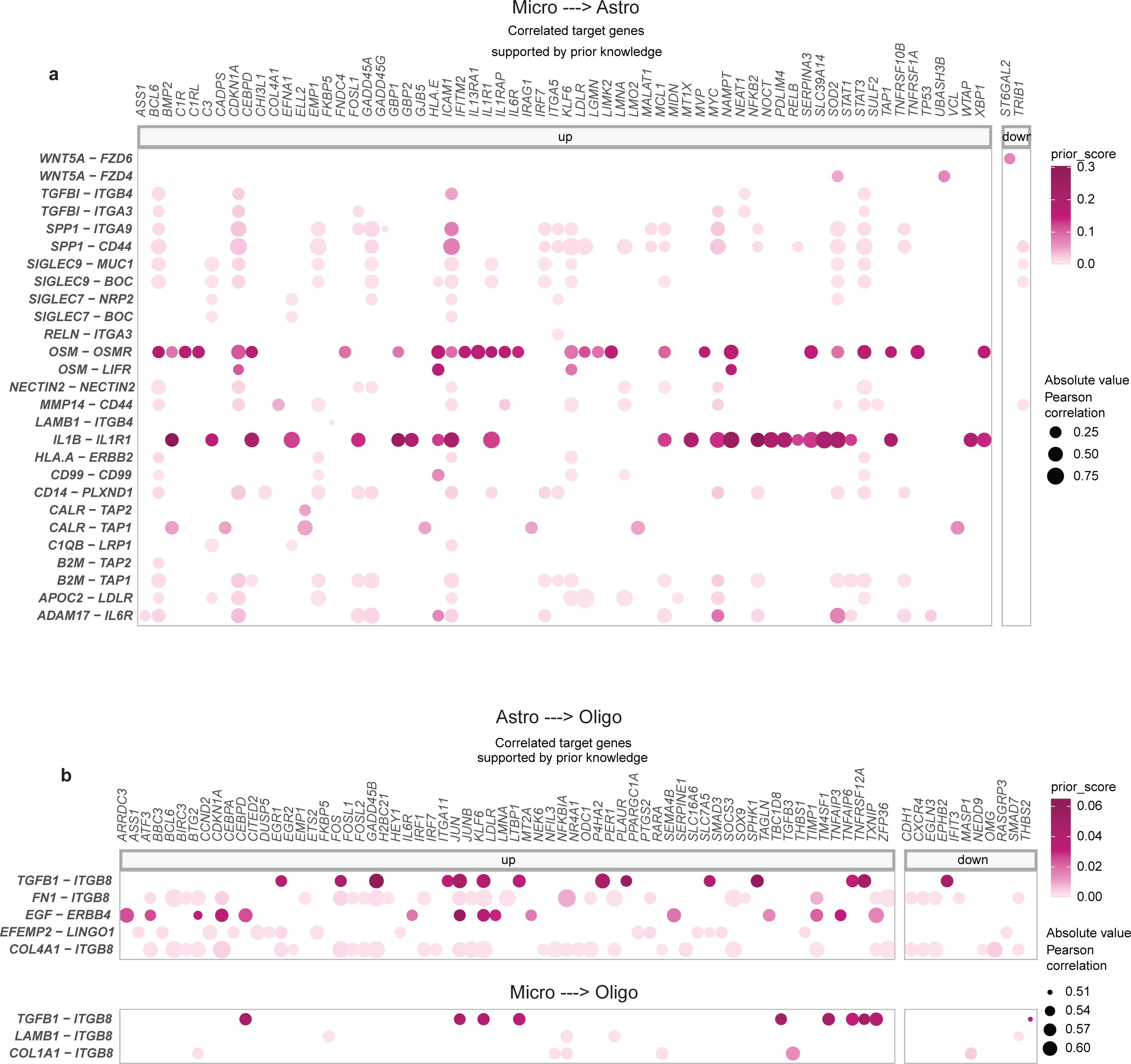
Downstream genes of AUD-associated ligand-receptor pairs. **a.** Microglia to astrocytes ligand-receptor pairs from MultiNicheNet cell-cell communication. All ligand-receptor pairs and target genes with high expression correlation (Spearman or Pearson > 0.50), having some prior knowledge to support their link (in the top 250 predicted target genes for the ligand, ‘prior score’ as predicted by MultiNicheNet), and being within the top 50 ligand-receptor pairs associated with AUD (as calculated by MultiNicheNet) are shown. Size of dots indicate Pearson correlation between expression of ligand-receptor pair and target gene. Color of dot indicates prior score for link between ligand-receptor and downstream gene. **b,** As **a**, with astrocytes to oligodendrocytes pairs (above) and microglia to oligodendrocyte pairs (below)

## Online Methods

### Sample Collection

The caudate from post-mortem brains of 183 donors were initially included in this study. Tissue was obtained from the New South Wales Brain Tissue Resource Centre (NSWBTRC), University of Sydney, Australia (https://sydney.edu.au/nsw-brain-tissue-resource-centre).^89^

### Genotype Data Processing and Imputation

NSWBTRC samples were genotyped using the UK Biobank Axiom® Array (ThermoFisher Scientific, Waltham, MA, U.S.A.). Before imputation, palindromic SNPs and SNPs with genotyping rate <95%, minor allele frequency (MAF) <1%, or Hardy-Weinberg equilibrium P-value < 1E-4 were excluded. Genotype data was imputed by using the TOPMed Imputation Server.^90^ Eagle v2^91^ was used to phase the genotypes and Minimac4 v1.2.4^90^ was used for imputation. Data from the Trans-Omics for Precision Medicine (TOPMed r3)^92^ was used as the reference genomes.

### Single-cell Multiome Assay

#### Nuclei Isolation for Single-cell Multiome

183 fresh-frozen post-mortem caudate brain samples were utilized in the assay. The 183 specimens were divided into 23 pools, with 8 in each pool. The donors in each pool were both condition (with or without AUD) and sex balanced. For each pool, around 20 mg tissue from each donor specimen was collected and combined into a sterilized 2 ml Dounce homogenizer. 2 ml chilled NP40 lysis buffer (10 mM Tris-HCl, pH 7.5, 10 mM NaCl, 3 mM MgCl2, 0.1% Nonidet P40 Substitute, 1mM DTT, 1 U/µl RNase inhibitor) was added to the Dounce homogenizer before the tissues were thawed. The tissues were homogenized 15x using pestle A, and 10x using pestle B, and were transferred into a centrifuge tube to incubate for 2 minutes on ice. After that, 2 ml wash buffer containing PBS, 1% BSA and 1 U/µl RNase inhibitor was added and mixed well. The lysed tissue was centrifuged at 500 rcf for 5 minutes at 4°C, then washed twice more with wash buffer and filtered through 70 µm and then 40 µm cell strainer separately. The pellet was resuspended in 2 ml wash buffer and mixed with 3.6 ml Sucrose Cushion Buffer I (nuclei PURE prep isolation kit, Sigma) containing 1 U/µl RNase inhibitor. 2 ml Sucrose Cushion Buffer I with 1 U/µl RNase inhibitor was added into one 15 ml Beckman Coulter centrifuge tube. After that, the 5.6 ml nuclei suspensions were gently added to the top of Sucrose Cushion Buffer I without mixing, and followed by centrifuging at 13,000 x rpm (30,000 rcf) (Beckman Coulter ultracentrifuge) with rotor SW40Ti for 45 minutes at 4°C. The purified nuclei pellet was washed by centrifuging at 300 rcf for 5 min at 4°C with wash buffer, and the washed nuclei pellets was resuspended in wash buffer to target ∼ 1000 nuclei/μl.

#### 10X Single-cell Multiome Library Preparation and Sequencing

Paired ATAC and gene expression libraries were generated following the Chromium Next GEM Single Cell Multiome ATAC + Gene Expression User Guide CG000338_RevB (10X Genomics, Inc). In brief, the isolated nuclei from a pool of samples were first incubated in a transposition mix. The single nuclei master mixture containing tagmented single nuclei suspension was loaded into two well of a Next GEM Chip J, along with the single cell multiome gel beads and partition oil. The chip was then loaded to the Chromium X Controller for GEM generation and barcoding. Barcoded transposed DNA and cDNA were amplified after GEMs being released. At each step, the quality of cDNA, ATAC library and cDNA library was examined by Bioanalyzer 2000. The final single indexed ATAC libraries and the dual indexed gene expression libraries were sequenced on an Illumina Novaseq 6000, with index reads of 10 bp + 24 bp, and 100 bp paired-end reads.

#### Cell Ranger ARC Analysis

Cell Ranger ARC (cellranger-arc-2.0.0, http://support.10xgenomics.com/) was utilized to process the raw sequence data derived from the single-cell multiome libraries. Both the ATAC and gene expression FASTQ files were processed with the cellranger-arc count algorithm. The reference refdata-cellranger-arc-GRCh38-2020-A-2.0.0 (10x Genomics) was used. The filtered gene-cell barcode matrices and fragment files were used for further analysis.

### Single-nuclei RNA-seq Assay

#### Nuclei Isolation for Single-nuclei RNA-seq

170 fresh-frozen post-mortem caudate brain samples (same individuals as in the multiome assay) were grouped into 17 pools, with 10 in each pool. The donors in each pool were both condition (with or without AUD) and sex balanced. The nuclei isolation for each pool is similar to the procedure described for the aforementioned single cell multiome assay. Zero point two (or One fifth) unit per microliter of RNase inhibitor was used in the buffers.

#### 10X HT Single-nuclei RNA-seq Library Preparation and Sequencing

The Chromium Next GEM single cell 3’ HT reagent kits v3.1 (user guide CG000416, 10X Genomics, Inc.) was used for the single-nuclei RNA-seq assay. The single nuclei suspension from a pool of 10 donor tissue samples were loaded into two wells of a Chromium Next GEM chip M to target 60,000 cell recovery per well. The chip was run on a Chromium X (10x Genomics). Single cell gel beads in emulsion containing barcoded oligonucleotides and reverse transcriptase reagents were generated. cDNA was synthesized and amplified following cell capture and cell lysis, The quality and quantity of cDNA and resulting libraries were examined by Bioanalyzer. The final libraries were sequenced on an Illumina NovaSeq 6000. 100-bp reads including cell barcode and UMI sequences and 100-bp RNA reads were generated.

#### Cell ranger Count Analysis

Cell Ranger Count (cellranger-count-7.0.1, http://support.10xgenomics.com/) was utilized to process the raw sequence data derived from the single-cell multiome libraries. The gene expression FASTQ files were processed with the cellranger count algorithm. The reference refdata-cellranger-GRCh38-2020-A (10x Genomics) was used. The filtered gene-cell barcode matrices and fragment files were used for further analysis.

### Demultiplexing

Cells from each of the 40 sequencing pools (23 from the single-cell multiome assay and 17 from the single-nuclei RNA-seq assay) were demultiplexed back into their samples of origin using the tool Demuxlet^93^ with default parameters, which uses genotype variant information for each sample to predict the sample of origin for each cell barcode, as well as identify doublet cells, artifactual libraries generated when two cells are captured in the same droplet during library preparation. Between 55% and 75% of cells from each pool were identified as singlets and assigned to a sample. The remaining cells (identified as doublets or ambiguous) were removed from further analysis.

After demultiplexing, seven samples from the HT assay and twenty-three samples from the multiome assay with either no genotype info available or less than 100 cell barcodes assigned were removed from all further analyses. Raw data from the barcodes from these samples are publicly available, see “Data Availability”, below. These samples – 163 from the HT experiments, and 161 from multiome – were used for all following processing and analyses until the filtering step detailed in ‘Sample Filtering’, below.

### Initial Quality Control

Unless specified differently, all following analysis was performed in R (version 4.3.1), predominantly utilizing the Seurat^94^ (v5) and Signac^95^ (v5) packages.

A Seurat object was created from the data from each pool from the HT assay using the gene expression count matrix from the Cell Ranger output (17 objects total). Cells with below 800 or above 11,250 genes, above 125,000 molecules, or above 10% mitochondrial RNA were removed from further analysis. These are commonly used quality control metrics to remove low-quality cells or multiplets. Each pool was then normalized using the scTransform() function in Seurat.

A Signac object, containing both RNA and ATAC-seq data, was created for each pool from the multiome assay from the hdf5 file from the Cell Ranger output (23 objects total). Cells with below 800 or above 20,000 genes, below 800 or above 500,000 detected RNA molecules, or above 20% mitochondrial RNA were removed from further analysis. An additional round of filtering was performed using the ATAC-seq data. The following cells were removed from further analysis:

- Cells with less than 100 or over 100,000 features;
- Less than 100 or over 1,000,000 counts;
- TSS enrichment less than 2;
- Nucleosome signal greater than 4;
- Percentage of reads in peaks less than 15%;
- Total number of fragments in peaks less than 800 or over 100,000;
- Ratio reads in genomic blacklist regions greater than 0.05 Between both assays, 1,307,323 unique barcodes passed all QC filters.

### RNA-seq Integration and Visualization

After the above quality control, all cells from the Seurat objects for each pool were integrated into the same Seurat object for visualization in the same 2D space. The atomic sketch integration method was used, a dictionary learning based procedure recently developed in Seurat for large datasets (see https://satijalab.org/seurat/articles/parsebio_sketch_integration). Briefly, 5,000 representative cells were selected from each pool (based on statistical leverage). Integration was performed on these sketched cells using the reference-based RPCAIntegration method. Then, each cell from each pool was placed in this integrated space as well using the ProjectIntegration function.

To visualize all cells in the same plot, we used functionality in the Seurat v5 and BPCells packages to convert each pool to an on-disk BPCells matrix.^96^ This allowed us to merge each object in a memory-efficient way. After merging, the function RunUMAP was run on the combined object for 2D visualization.

### Cell Type Annotation

The 1,307,323 cells were divided into 49 clusters using the FindNeighbors and FindClusters functions in Seurat. Cell clusters were annotated into known striatal cell types based on expression levels of a combination of marker genes curated from established studies (Figure 2B).^32,33,41,62,97^

### ATAC-seq Integration, Visualization

Each of the 23 Signac objects were processed using the standard ATAC-seq procedure in Signac – FindTopFeatures, RunTFIDF, RunSVD – and all objects were then integrated using the IntegrateEmbeddings command.

Cell type labels were transferred to the ATAC-seq object using the assignments for each barcode determined from the snRNA-seq data. Barcodes without a matching QC-passed snRNA-seq barcode were excluded, leaving 250,537 cells from 159 samples. UMAP visualization was calculated with the RunUMAP command, using the integrated_lsi reduction determined in the previous integration step.

### Sample Filtering

Following the above analyses, 20 samples with a proportion of glutamatergic neurons greater than 10% were removed from both the processed RNA-seq and ATAC-seq data, because such a cell-type composition indicates potential contamination with non-caudate tissue, leaving 143 samples for the following downstream analyses.

### Cell Subtype/Substate Annotation and Testing

Microglia and astrocyte clusters were further divided into four and two subclusters, respectively, by performing another iteration of FindNeighbors and FindClusters on these individual clusters, using cells from the 143 samples. Subcluster-specific genes were determined by using the “roc” test within the FindMarkers function in Seurat (see https://satijalab.org/seurat/reference/findmarkers). The top 50 genes (based on myAUC statistic) were used as input into g:Profiler for each subcluster to determine enriched biological pathways specific to that subcluster.

For testing for a difference in the proportion of cell states in individuals with AUD, the mean proportion of cells in each cluster were calculated for each sample, and an ANOVA was performed to determine if the mean proportion significantly changed in samples from individuals with AUD as compared to those without. Age, sex, and ethnic origin were used as covariates. For this test, we removed samples with fewer than 50 cells of the cell type being tested, as these samples contain very few cells of each subcluster and thus their mean is more unreliable.

### Differential Expression Analysis

#### RNA Pseudobulk Samples Creation

Due to the sparsity of single-cell data, differential expression methods designed to be run on the single-cell level often lack high statistical power. To account for this challenge, we utilized a pseudobulk approach. To create the pseudobulk data, for each cell type, the gene expression matrices of each cell of that cell type were combined (summed) by sample ID. Samples were removed on a cell type-specific basis if the sample contained less than 50 cells of that cell type. See Supplementary Table 7 for a summary of the number of pseudobulk samples created for each cell type. Due to a low number of samples (less than 10 individuals with AUD and 10 without) meeting the >=50 cell criteria, differential gene expression analysis was not performed for cholinergic interneurons, vascular smooth muscle cells, CCK interneurons, and macrophages. These pseudobulk samples were used for the differential expression analysis, below.

#### Differential Gene Expression Analysis

Differential gene expression analysis between samples from individuals with and without AUD was performed for each cell type, as well as the two subclusters of astrocytes and four subclusters of microglia, using DESeq2,^98^ a statistical package designed for bulk RNA-seq data, with the default parameters. Briefly, the tool estimates the variance of gene expression and then fits a negative binomial distribution to each gene, which accounts for the over-dispersion of RNA sequencing data, which can result in more accurate p-values. Sex, age (as a continuous variable), and ethnic origin were included as covariates in the models. Genes with p values of less than 0.2 (corrected for multiple-hypothesis testing using the Benjamini-Hochberg method) were deemed significant.

### Gene Set Enrichment Analysis

Gene set enrichment analysis was performed for each cell type that underwent differential expression analysis, using the fgsea R package,^99^ which uses a preranked list of genes to determine gene sets that are enriched based on the gene rankings. In this case, the log_2_ fold changes from the differential expression analysis were used as the ranks, and pathways from the Reactome database were used as gene sets. See Supplementary Table 9 for full fgsea results for each cell type. For visualization, the top 30 enriched pathways (based on smallest Benjamini-Hochberg-adjusted *p* values) in each cell type were selected and hierarchically clustered based on number of genes shared between the pathways. Clustered pathways were then manually labeled into 25 groups.

### Creation of Cell Type-specific ATAC-seq Profiles

CoveragePlot function in Signac was used for visualization of ATAC-seq signal for marker genes.

Peak calling was performed separately for cells from each of the 16 cell types (excluding glutamatergic neurons) using the CallPeaks function in Signac with default parameters. The function uses MACS2^100^ for peak calling.

For comparing similarity of peaks called between cell types, the Jaccard index was used, defined here as the number of peaks in one cell type overlapping a peak in the other cell type, divided by the union of the peaks in both cell types.

All cells from the integrated ATAC-seq object, from the 143 samples determined after the ‘Sample Filtering’ step, above, were used for the procedures in this section.

### Differential Chromatin Accessibility Analysis

#### ATAC Pseudobulk Samples Creation

In the same way as the RNA-seq data, pseudobulk chromatin accessibility data was created for the ATAC-seq data: For each cell type, the ATAC-seq counts matrices of each cell of that cell type were combined (summed) by sample ID, for the 143 samples. Samples were removed on a cell type-specific basis if the sample contained less than 50 cells of that cell type. See Supplementary Table 12 for a summary of the number of pseudobulk samples created for each cell type. Due to a low number of samples meeting the >=50 cell criteria, differential accessibility analysis was not performed for cholinergic interneurons, vSMCs, CCK interneurons, macrophages, ependymal cells, LTS interneurons, or endothelial cells.

#### Differential Accessibility Analysis

Differential chromatin accessibility analysis between individuals with and without AUD was performed for each cell type using DESeq2 with the default parameters. Sex, age, and ethnic origin were included as covariates in the models. Genes with p values of less than 0.2 (corrected for multiple-hypothesis testing using the Benjamini-Hochberg method) were deemed significant. Regions residing in promoter regions of genes was determined using R package ChIPSeeker.^101^ Namely, the function annotatePeak() was used, with parameters: TxDb = TxDb.Hsapiens.UCSC.hg38.knownGene, annoDb=“org.Hs.eg.db”, and tssRegion = c(-1000, 1000).

#### Comparison of Differentially Accessible Genes and Differentially Expressed Genes

To calculate the association between gene expression and chromatin accessibility differences for each gene, we assigned to each gene with at least 1 DAR in the promoter region the log_2_ fold change of the DAR with the highest ATAC signal, as well as the log_2_ fold change of the gene’s expression.

For the GSEA analyses, the log_2_ fold changes from the differential expression analysis were used as the ranks, and genes with at least 1 DAR in the promoter region were used as the gene sets, separated into genes with positive log_2_ fold changes, and those with negative log_2_ fold changes.

### Identifying Genes Containing Significant GWAS Loci

The two GWAS studies (Saunders, et al.^3^ and Zhou et al.^5^) utilized in this study defined a locus as the region including all variants in linkage disequilibrium of r2> 0.1. For the present study, all 1,307 loci associated with the drinks per week (DrnkWk) phenotype from Saunders et al. were selected. To find genes overlapping with these regions, gene annotations from UCSC Genome Browser for the hg38 genome assembly (implemented in the TxDb.Hsapiens.UCSC.hg38.knownGene R package^102^) were used. This identified 3,406 genes. A comparable process was used to select genes overlapping loci from the Zhou et al. study. All 75 loci associated with the problematic alcohol use phenotype were selected, which overlapped with 750 known genes from the UCSC annotations.

### Variant Calling

Integrated snATAC-seq data were split into single bam files for each of the 143 individuals using sinto (https://timoast.github.io/sinto/). Duplicated bam files for the same sample were merged together with samtools.^103^ For variant calling, the Sentieon germline variant calling pipeline^104^ was used, namely:

Removal of duplicate RNA molecules;

> Recalibration of base quality score using GATK’s Base Quality Score Recalibration;
>
> Variant calling was performed for each sample using the Sentieon’s Haplotyper algorithm;
>
> Joint variant calling was performed using the DNAseq algorithm;

Variants were recalibrated using GATK’s Variant Quality Score Recalibration^105^ algorithm; variants not passing the recalibration test were filtered out for further analyses.

### eQTL Analysis

Expression quantitative trait loci (eQTL) analysis was performed for all SNPs in each cell type within 100,000 bases of a differentially expressed gene overlapping a GWAS loci and within an ATAC-seq peak from that cell type. tensorQTL^106^ was used to perform the analysis using the map_cis function with default parameters. Variants with minor allele frequency < 0.05 were excluded from the analysis. Sex, age, and ethnic origin were used as covariates.

### Gene Regulatory Mechanisms Prediction

#### chromVAR^107^

Prediction of motif activities for each cell was performed using the RunChromVAR command in Signac. Briefly, chromVAR identifies motifs associated with variability in chromatin accessibility between cells^107^). Differential testing on the chromVAR z-score was performed using the FindMarkers function, setting mean.fxn=rowMeans and fc.name=“avg_diff”, so that the fold-change represents the average difference in z-score between the groups.

To make these differential activity motif results more robust, we utilized a pseudobulk approach: averaging per-cell motif scores for all cells within a sample of a given cell type, taking the log, and then using an ANOVA – with sex, age, and ethnic origin as covariates – to test for differences between those with and without AUD. All samples used for differential accessibility testing (see ‘ATAC Pseudobulk Samples Creation’) were used for this analysis.

#### Gene Regulatory Network Inference

To build the cell population gene regulatory network, we used LINGER, as described in Yuan & Duren, 2024.^43^ We generated pseudobulk-level expression and chromatin accessibility data for each donor and each cell type. Here we used the union set of peaks (described above) for ATAC-seq data. as well as a covariate matrix, with sex, age, and ethnic origin as covariates to the model. All samples used for differential accessibility testing (see ‘ATAC Pseudobulk Samples Creation’) were used for this analysis.

#### Trans-regulatory module detection by matrix factorization

To detect key TF-TG subnetworks (modules) from the cell population TF–TG trans-regulation, we used non-negative matrix factorization (NMF). Before matrix factorization, We normalized the trans-regulatory potential matrix by standardizing each row (TF) and each column (TG) independently. The standardization of each TF ensures that for each TF, the average regulatory potential across TGs becomes zero, and the variation in regulatory potential across genes has a standard deviation of one. The same normalization was applied to each TG, so that the effect of the regulator side was also normalized. We took the average of these two standardized matrices and set the negative values to zeros, which was used for downstream analysis. Next, we performed NMF on the preprocessed matrix to decompose it into two non-negative matrices, *W* and *H*, representing module membership of TGs and TFs, respectively. *W* is a *m* by *k* module weight matrix for TF, representing module weight of TF, where m is the number of TF. *H* is a *k* by *n* matrix, representing module weight for TGs, where n is the number of TG. To assign TF and TG into specific modules, we normalized the module weight matrix to equal sum for different modules. For each gene, we converted the normalized weight matrix into proportions by dividing the sum of weights across modules. We sorted genes based on their highest proportion across all modules to select the top 10% of genes and assign them to modules for which the gene has the largest score.

This procedure was applied to TF module W matrix and the TG module *H* matrix. Here, we identified 10 trans-regulatory modules.

To uncover AUD-associated regulatory programs in each cell type, we performed differential module expression analysis. We first pre-processed the pseudo-bulk gene expression count matrix by: (1) normalizing for cell depth, (2) log-transforming, and (3) z-scoring expression across all donors. We then estimated module activity in each donor as the mean expression of module genes. Using a two-tailed two-sample t-test, we identified differentially active modules between individuals with and without AUD.

#### Identification of cis and trans driver TFs

We identified *cis* and *trans* driver TFs underlying epigenetic and transcriptome changes between individuals with and without AUD using a linear regression model, *Y* = *Aβ* + *β*_0_ + *ε*. For transcriptome drivers, the regression model predicted the log transformation of the gene expression fold change between individuals with and without AUD (Y) from cell type-specific TF-TG trans-regulation (A). For epigenetic drivers, the model predicted chromatin accessibility changes (Y) from the cell type-specific TF-regulatory element cis-regulation (A). Significant TFs from each model indicated TFs driving differential expression and chromatin states between conditions through direct epigenetic or transcriptome regulation.

### Cell-cell Communication Analysis

To analyze cell-cell communication differences in individuals with AUD, we used MultiNicheNet,^55^ an R package for differential cell-cell communication analysis using single-cell data with multi-sample, multi-condition designs. All samples used for differential expression testing (see ‘RNA Pseudobulk Samples Creation’) were used for this analysis. User-set parameters were set as the following:

○ MultiNicheNet’s analysis uses a pseudobulk approach, and the minimum number of cells per cell type per sample was set to 10, the recommended default;
○ Sex, ethnic origin, and age were used as covariates in the design;
○ For a differentially expressed gene to be further considered when calculating ligand activity, we choose for a minimum log_2_ fold change of 0.50, maximum adjusted *p*-value of 0.2, and minimum fraction of expression of 0.05;
○ For the NicheNet ligand-target inference, the top 250 predicted target genes were considered;
○ The weights of the prioritization of expression, differential expression and NicheNet activity information was set to the recommended defaults (see https://github.com/saeyslab/multinichenetr);
○ Sender cell types were defined as astrocytes, microglia, and oligodendrocytes;
○ Receiver cell types were defined as astrocytes, microglia, and oligodendrocytes;

Visualization of top 5 ligand-receptor pairs in individuals with AUD, based on scaled ligand activity score, was created using the make_circos_group_comparison function.

## Data Availability

The GWAS datasets utilized in this study were obtained from Saunders, et al.^3,5^ – summary statistics of which can be found at https://conservancy.umn.edu/handle/11299/241912 – and Zhou, et al.^5^ – of which full summary-level information can be found at https://medicine.yale.edu/lab/gelernter/stats/ and dbGaP (accession number phs001672).

The data generated here, including raw sequencing data in the form of BAM files and processed data in the form of Seurat RDS objects will be available at time of publication.

## Code Availability

No custom computational packages or extensive computer code were developed in this study. However, R or Python scripts used to utilize existing packages are available from the corresponding author upon request.

## Notes

### Competing Interest Statement

The authors have declared no competing interest.

### Summary of Updates

Revised abstract and discussion sections. New results added to gene regulatory network section. Author affiliation updated. Minor edits to figures.

